# The effects of direct brain stimulation in humans depend on frequency, amplitude, and white-matter proximity

**DOI:** 10.1101/746834

**Authors:** Uma R. Mohan, Andrew J. Watrous, Jonathan F. Miller, Bradley C. Lega, Michael R. Sperling, Gregory A. Worrell, Robert E. Gross, Kareem A. Zaghloul, Barbara C. Jobst, Kathryn A. Davis, Sameer A. Sheth, Joel M. Stein, Sandhitsu R. Das, Richard Gorniak, Paul A. Wanda, Daniel S. Rizzuto, Michael J. Kahana, Joshua Jacobs

## Abstract

Researchers have used direct electrical brain stimulation to treat a range of neurological and psychiatric disorders. However, for brain stimulation to be maximally effective, clinicians and researchers should optimize stimulation parameters according to desired outcomes. To examine how different kinds of stimulation affect human brain activity, we compared the changes in neuronal activity that resulted from stimulation at a range of frequencies, amplitudes, and locations with direct human brain recordings. We recorded human brain activity directly with electrodes that were implanted in widespread regions across 106 neurosurgical epilepsy patients while systematically stimulating across a range of parameters and locations. Overall, stimulation most often had an inhibitory effect on neuronal activity, consistent with earlier work. When stimulation excited neuronal activity, it most often occurred from high-frequency stimulation. These effects were modulated by the location of the stimulating electrode, with stimulation sites near white matter more likely to cause excitation and sites near gray matter more likely to inhibit neuronal activity. By characterizing how different stimulation parameters produced specific neuronal activity patterns on a large scale, our results help guide clinicians and researchers when designing stimulation protocols to cause precisely targeted changes in human brain activity.

## Introduction

Direct electrical stimulation shows potential as a treatment for a variety of neurological conditions and as a tool for studying neuropsychiatric disorders and cognition. However, we do not yet have a detailed understanding of the widespread neuronal effects that result from different types of stimulation. The goal of our study was to examine this issue by characterizing at a large scale how different types of brain stimulation modulate directly recorded human neuronal activity.

For years, direct electrical stimulation has been used to effectively treat motor disorders, such as Parkinson’s Disease, essential tremor, dystonia, and epileptic seizures [Benabid et al., 1987, Lang and Lozano, 1998a,b, Koller et al., 1997, Kumar et al., 1999, Coubes et al., 2000, Yianni et al., 2003, Fisher et al., 2010, Fisher and Velasco, 2014]. In the past two decades, researchers have extended stimulation protocols from motor disorders to better understand and modulate brain circuits of neuropsychiatric and cognitive disorders, such as major depression [Mayberg, 1997], obsessive compulsive disorder [Nuttin et al., 2003], anorexia nervosa [Lipsman et al., 2017], addiction [Kuhn et al., 2007, Levy et al., 2007], schizophrenia [Kuhn et al., 2011, Bakay, 2009], and Alzheimer’s disease [Kuhn et al., 2015, Lozano et al., 2016]. While direct electrical stimulation holds potential to treat patients with neurological disorders who cannot be treated pharmacologically, understanding more fully how different stimulation parameters differentially affect neuronal activity is important for optimizing such therapies.

Researchers and clinicians have found that stimulation produces a wide range of behavioral effects. Cortical stimulation was first linked to memory in Wilder Penfield’s pioneering studies where stimulating an awake patient’s temporal lobe caused them to spontaneously recall old memories [Penfield and Perot, 1963]. Penfield’s subsequent work showed that the particular location that was stimulated greatly affected the way in which patients re-experienced old memories. Following this, many studies applied direct electrical stimulation to the temporal lobe using a variety of stimulation parameters. The results from these studies were wide-ranging, emphasizing the complexity of precisely modulating human neuronal activity with stimulation [Selimbeyoglu and Parvizi, 2010, Borchers et al., 2012, Suthana and Fried, 2014, Ezzyat and Rizzuto, 2018]. Some studies showed that stimulation impaired recall of complex scenes [Halgren et al., 1985], subsequent item recognition [Coleshill et al., 2004], spatial, and verbal memory recall [Jacobs et al., 2016, Lacruz et al., 2010]. However, a number of studies have also shown improvements to verbal, visual, and spatial memory [Suthana et al., 2012, Fell et al., 2013, Miller et al., 2015, Ezzyat et al., 2017]. Studies using brain stimulation to treat other neurological diseases also found inconsistent cognitive effects [Gutman et al., 2009, Mayberg et al., 2005, Lang and Lozano, 1998a,b]. There were substantial variations in stimulation protocols between these studies, include stimulation location, frequency, duration, amplitude, pulse pattern (continuous or intermittent), and timing. To explain why these studies found such diverse behavioral and cognitive effects from stimulation, it is helpful to understand the physiology of how different kinds of stimulation alter underlying neuronal activity.

Earlier studies showed that stimulation can cause both excitatory and inhibitory effects on local and connected regions. Yet, within the realm of treating Parkinson’s Disease with deep brain stimulation (DBS) where clinical outcomes are well established, the electrophysiology of stimulation is unclear. While some studies demonstrate that stimulation causes inhibition [Limousin et al., 1995, Welter et al., 2004, Boraud et al., 1996, Dostrovsky et al., 2000], other studies show excitation after stimulating at different frequencies and locations [Anderson et al., 2003, Hashimoto et al., 2003, Maurice et al., 2003, Windels et al., 2000, Johnson et al., 2008]. There is evidence that the location of a stimulation electrode also has an important role in dictating the outcome of stimulation, with white- and gray-matter stimulation sites causing different effects [Histed et al., 2009, 2013, Nowak and Bullier, 1998a,b]. Further, Logothetis et al. [2010] show evidence in monkeys that specific patterns of stimulation can simultaneously induce inhibitory both excitatory effects in different affected regions. These findings, which illustrate the diverse range of electrophysiological effects that result from brain stimulation, demonstrate the challenge in designing brain stimulation protocols to alter brain activity in targeted ways that achieve desired behavioral outcomes.

The goal of our study was to comprehensively evaluate the effects of different types of stimulation on neuronal activity across the human brain. To examine changes in neuronal activity due to stimulation, we collected and analyzed direct brain recordings from 106 neurosurgical patients who underwent an extensive stimulation “parameter search” paradigm involving a range of stimulation frequencies and amplitudes at different cortical surface and depth locations. We then measured how different stimulation parameters correlated with the directional changes in neuronal activity that resulted from stimulation. Because we sought to understand the effects of stimulation on the mean activity across neuronal populations, we measured high-frequency broadband power, which provides an estimate of the mean rate of local neuronal spiking activity [Manning et al., 2009, Watson et al., 2018]. Our results provide a more comprehensive study of the direct electrical stimulation parameter space than any prior human study. We find that the neuronal effects of stimulation are highly parameter dependent. Specifically, the prevalence of excitation and inhibition are modulated by the frequency and amplitude of stimulation and by the distance of the stimulation site to white-matter tracts. These results provide guidance for clinicians and researchers to more optimally craft stimulation parameters according to the desired types of changes to ongoing brain activity.

## Methods

### Participants

The 106 patients in our study were surgically implanted with depth, surface grid, and/or surface strips of electrodes for the purpose of identifying epileptic regions. The patients’ clinical teams determined electrode placement to best monitor each patient’s epilepsy. We conducted these procedures at eight hospitals: Thomas Jefferson University Hospital (Philadelphia, PA); University of Texas Southwestern Medical Center (Dallas, TX); Emory University Hospital (Atlanta, GA); Dartmouth–Hitchcock Medical Center (Lebanon, NH); Hospital of the University of Pennsylvania (Philadelphia, PA); Mayo Clinic (Rochester, MN); National Institutes of Health (Bethesda, MD); and Columbia University Hospital (New York, NY). Following institutional review board protocols at each hospital, all participating patients provided informed consent.

### Stimulation Paradigm

This stimulation “parameter search” paradigm was part of a larger project aimed to enhance episodic and spatial memory using direct electrical stimulation [Jacobs et al., 2016, Ezzyat et al., 2017, 2018]. Blackrock Microsystems provided neural stimulation equipment for these protocols. As part of this larger project, subjects participated in this paradigm to characterize the brain-wide effects of applying electrical stimulation at different sites with varying frequencies and amplitudes. During each session of this stimulation procedure, we instructed subjects to sit quietly and rest with eyes open as we applied various types of stimulation and measured neuronal activity. The main goal in applying stimulation across frequencies, amplitudes, and sites was to identify specific stimulation locations and parameters that would enhance performance in a subsequent memory task [Ezzyat et al., 2017]. Therefore, stimulation was often applied in MTL and lateral temporal lobe locations based on their functional relevance for memory [Eichenbaum, 2000, Ojemann et al., 1989], as well as other areas (Table S2).

A clinical neurologist oversaw all stimulation sessions. We performed a separate amplitude screening procedure before beginning stimulation for each target site. In the screening procedure, each site was progressively stimulated for 0.5 s at each tested frequency, beginning at 0.5 mA, in steps of 0.5 mA, up to a maximum of 1.5 mA for depth electrodes or 3 mA for surface electrodes. A neurologist monitored visually for afterdischarges throughout this process. We then logged for each site the maximum current that could be applied without causing afterdischarges.

Then, in the main stimulation protocol for each site, we applied bipolar stimulation across neighboring anode and cathode electrodes using 300-*µ*s charge-balanced biphasic rectangular pulses. For each site, we stimulated at frequencies of 10, 25, 50, 100, or 200 Hz, with amplitudes from 0.25 mA up to the site’s determined maximum in steps of 0.25 mA, as well as 0.125 mA. Each stimulation trial was applied for 500 ms, with a random delay of 2750–3500 ms (uniformly distributed) between the offset and onset of consecutive stimulation trials. Within each *∼*25-minute session that targeted one stimulation site, we randomly ordered the stimulation trials with different frequencies and amplitudes for each site. Each targeted electrode received 24 stimulation trials for each combination of frequency and amplitude. Some subjects participated in a version of this procedure that also included sham trials without stimulation. Individual subjects participated in this stimulation protocol for between 1 and 9 individual sites (mean = 2.8 sites). Overall, we collected a total of 354 sessions, stimulating at 319 distinct sites from 106 subjects. Following artifact rejection (see below), we included in our data analyses 292 sessions over 263 stimulation sites from 94 subjects while recording simultaneous neuronal activity from 10,310 bipolar electrode pairs.

### Electrocorticographic recordings and referencing

To measure the electrophysiological effects of stimulation, throughout stimulation we recorded neuronal activity at 500, 1000, or 1600 Hz using a clinical intracranial electroencephalographic (iEEG) recording system at each hospital (Nihon Kohden EEG-1200, Natus XLTek EMU 128, Natus Quantum EEG, or Grass Aura-LTM64 systems). We referenced each electrode’s signal to a common contact placed intracranially, on the scalp, or mastoid process. To reduce non-physiological artifacts, we used bipolar referencing, computed as the voltage difference between pairs of adjacent electrodes. The location of each bipolar pair was taken as the midpoint between the two physical electrodes. We further filtered electrical line noise using a 57–63-Hz Butterworth notch filter.

### Anatomical localization

We determined the location of each electrode by co-registering a postsurgical CT scan to T1 and T2 weighted structural MRIs taken prior to implantation. We determined electrode localization in cortical regions by co-registration of the post-implantation CT, corrected for post-operative brain shift, with Freesurfer’s automated cortical parcellation based on the Desikan-Killiany brain atlas [Desikan et al., 2006]. We based localization to medial temporal lobe (MTL) structures on MTL segmentation using Automatic Segmentation of Hippocampal Subfields (ASHS) [Yushkevich et al., 2015].

### Artifact Rejection

Applying electrical stimulation can cause the appearance of non-physiological signals in iEEG recordings that may manifest as complete amplifier saturation as well as overall shifts in signal amplitude, such as rise, decay, or deflection following stimulation before returning to baseline (Fig. S2). These non-physiological changes could impair our ability to accurately measure true physiological signals related to stimulation. Therefore, to minimize the impact of artifacts on our results, we excluded from our analyses any recording electrodes and trials that showed post-stimulation artifacts. We implemented a detection algorithm to identify channels that are prone to complete signal saturation as well as gradual artifact following stimulation. Following earlier methods [Solomon et al., 2018], we compared the average voltage of the signal from *−*500 to *−*100 ms prior to stimulation onset and from 100 to 500 ms after stimulation offset. To include data from as many recording electrodes as possible, we took a two-phase approach to exclude artifacts on the single-trial level as well as on an electrode level. To identify artifacts, we employed Grubb’s outlier test to classify the trials that exhibited large non-physiological changes in voltage. Specifically, we excluded the data of any trials that showed a change in voltage between the pre- and post-stimulation intervals that was greater than 2 standard deviations of the corresponding mean voltage changes for matching sham trials for that electrode (Fig. S2). We excluded any electrodes completely that showed artifacts on more than half of all trials for a particular combination of parameters. Some stimulation sites were especially conducive to spreading artifacts across recording electrodes, and thus we excluded stimulation sites that caused artifacts on over half of all recording electrodes. Overall, we excluded 56 stimulation sites, an average of 10% of bipolar recording electrodes, and 12% of stimulation trials on remaining contacts (see Table S3).

### Spectral Power Analysis

To measure the effect of stimulation on mean neuronal firing rates, we extracted the high-frequency activity (HFA) signal from each iEEG recording, as this signal has been shown to provide a reliable measure of mean neuronal activity [Manning et al., 2009, Miller et al., 2009]. We measured HFA power in our data by calculating power spectra post-(200 to 700 ms after stimulation offset, defined as the last pulse of the stimulation trial) and pre-(*−*600 to *−*100 ms before stimulation onset) stimulation at 12 log-spaced frequencies between 30 and 100 Hz using multi-tapers, which provide better resolution at high frequencies [Mitra and Pesaran, 1999]. We allowed this buffer of 100 ms before and after stimulation to prevent any impact of stimulation artifacts on our results. In order to detect sites where activity resets to a specific level following stimulation, we calculated the variance of HFA power values before and after stimulation across all trials with the same combination of stimulation site, frequency, and amplitude. If variances are unequal and post-stimulation variance was less than the pre-stimulation variance, we categorized the site as showing “resetting.”

### Linear Mixed-Effects Model

We used a linear mixed-effects (LME) model to analyze the effects of stimulation on neuronal activity and identify how the prevalence of these effects vary with parameters. An LME model is a type of regression model that models the variation of a dependent variable as a function of both fixed and random effects. An LME model may be implemented in a group-based way that can account for repeated measurements from one sample [Baayen et al., 2008]. This feature is important for our study because our dataset included possibly correlated measurements, as we tested the effects of different parameters at the same stimulation site. Additionally, the LME model is useful for this dataset because it can account for uneven sampling across groups and conditions, which also occurred when separate sites were stimulated with different sets of frequencies and amplitudes.

To apply the LME model to our data, we used three random factors: frequency (up to 5 possible values per site), amplitude (usually 3 per site), and distance from the stimulation site (in Talairach units). We defined the direction of HFA change as a fixed factor because increases and decreases were the only changes of interest compared. For each stimulation site, the model fits either a random or fixed intercept and slope for each factor. Then the data across sites are combined to provide a summary coefficient for the factor that indicates the mean effect over all stimulation sites (groups) based on the normal distribution [Bates et al., 2014]. To compare the effects of stimulating distinct groups of electrodes, such as surface versus depth electrodes or white versus gray matter stimulation, we used a two-way ANOVA.

### Seizure-onset zones

Clinical teams at each hospital provided information about electrodes identified as in seizure onset zones. To verify that our results were not directly related to abnormal brain tissue, we performed the population analyses of the effects of stimulation frequency and amplitude for the sets of stimulation sites (n = 98) that were located in seizure onset zones (Fig. 2). All main frequency- and amplitude-related effects continued to be significant in this restricted analysis, confirming our main results.

**Figure 1:**
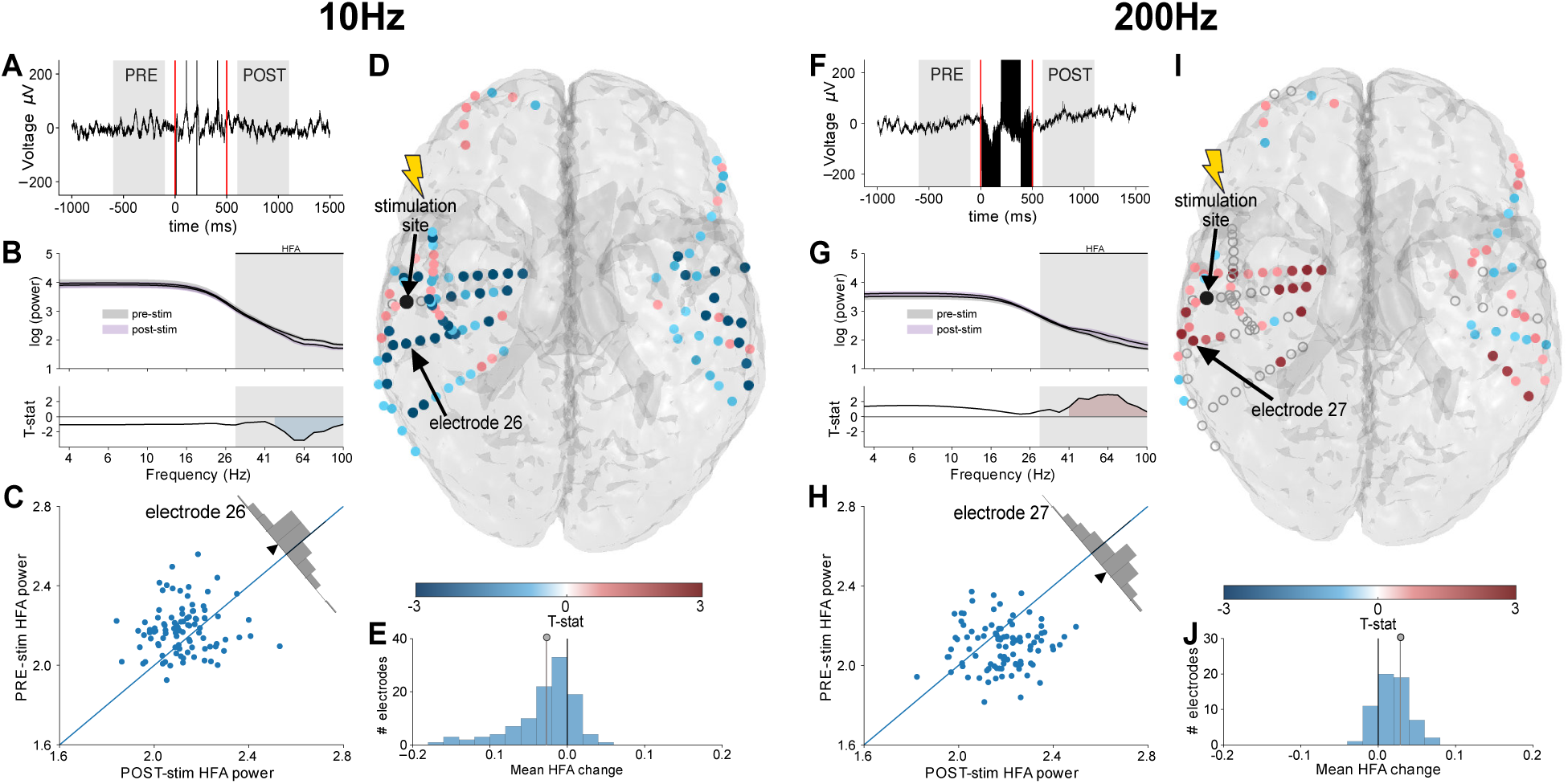
Effects of low- and high-frequency stimulation on HFA power. Left panels (A–E) indicate effects of 10-Hz stimulation and right panels (F–J) indicate 200-Hz stimulation, all in Patient 195. Stimulation was applied at the same site and amplitude (1 mA) for all panels. **(A)** Raw signal recorded on example electrode 26 on one trial. Shading indicates the 500-ms time periods before and after each stimulation trial during which we measured HFA power. Red lines denote stimulation onset and offset. **(B)** Top panel shows log-transformed mean power spectra from recording electrode 26 for the pre- and post-stimulation intervals across the 96 stimulation trials at 10 Hz and 1 mA. Gray shading indicates the HFA band (30–100 Hz). Bottom panel show t statistic of the difference between pre- and post-stimulation (POST-PRE) power at each frequency. Blue shading indicates significant differences at p<0.05. **(C)** The distribution of pre- and post-stimulation HFA power across individual trials for electrode 26. **(D)** Brain map showing the mean HFA responses to 10-Hz stimulation across all recording electrodes. The stimulation site is indicated in black and color indicates the t statistic of the change in HFA power at each recording electrode. Recording electrodes excluded due to artifact indicated by an open gray circle. **(E)** The distribution across electrodes, of the mean HFA power change in response to 10-Hz stimulation. Each value in this plot represents one electrode’s mean HFA power change from stimulation (POST-PRE). **(F–J)** Plots follow format from panels A–E except for 200-Hz stimulation with example data from recording electrode 27.

**Figure 2:**
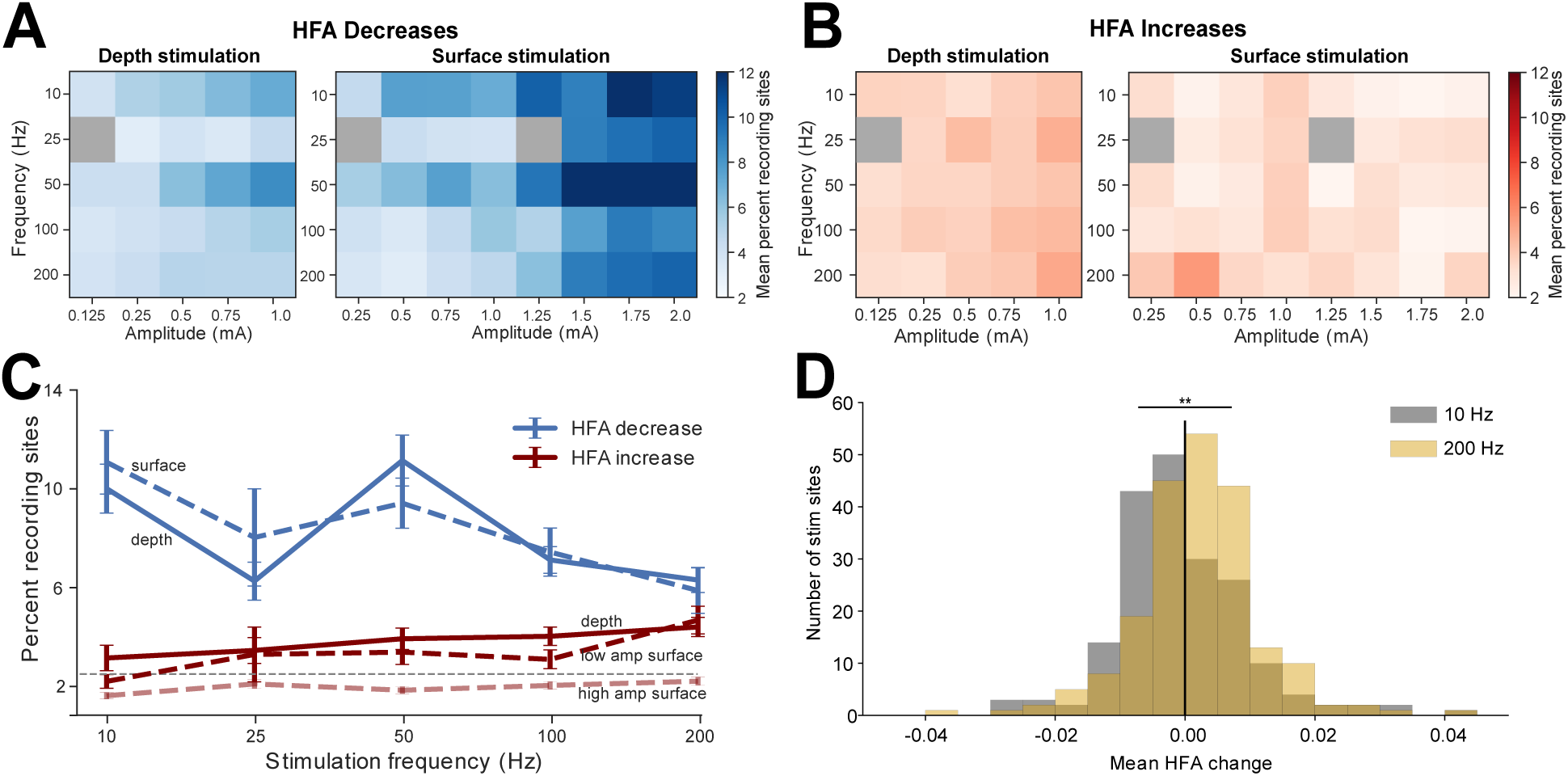
Population analysis of the frequency- and amplitude-dependence of HFA changes from stimulation. **(A)** Percent of recording electrodes showing significant HFA decreases for each combination of stimulation frequency and amplitude, separately computed for depth (left) and surface (right) stimulation. **(B)** Percent of recording electrodes showing significant HFA increases for each combination of stimulation parameters. **(C)** Prevalence of recording electrodes showing significant HFA increases and decreases for each stimulation frequency. Calculations were performed separately across depth and surface stimulation sites. Data in this plot included 1 mA stimulation for both effects of depth stimulation and surface HFA decreases; surface HFA in-creases calculations separately measured for currents ≥0.75 mA and <0.75 mA. Error bars: ±1 SEM. Gray dashed line indicates percent of electrodes increasing or decreasing by chance (2.5%). **(D)** Histogram of the mean HFA change for each stimulation site, separately computed for high and low-frequency stimulation; ** denotes a significant difference (z = *−*3.81*, p* = 0.0001*, rank-sum test)*.

### White matter categorization

We categorized each stimulation site as either being in/near white matter or in gray matter to determine the impact of white matter on the effects of stimulation. We estimated the white matter near each stimulation site by counting the number of white matter vertices within 3 Talairach units of the midpoint of the stimulation anode and cathode. We used Freesurfer white matter segmentation of patients’ T1 MRI scan to determine white matter vertex locations [Solomon et al., 2018]. We then categorized stimulation sites as near white matter or in gray matter by splitting the number of white matter vertices surrounding stimulation sites along the median of the distribution.

### Data Availability

Raw electrophysiogical data used in this study are available at http://memory.psych.upenn.edu/Electrophysiological_Data.

## Results

The goal of our study was to characterize the effects of different types of direct electrical brain stimulation on ongoing neuronal activity in humans. Here, we recorded intracranial electroencephalographic (iEEG) activity from widespread electrodes while delivering electrical stimulation at different locations, frequencies, and amplitudes as patients rested quietly. To assess the effect of stimulation on neuronal activity, we measured the amplitude of signals in the high-frequency-activity (HFA) range (30–100 Hz), which is an iEEG signal that correlates with the mean level of spiking activity across a local neuronal population [Manning et al., 2009, Watson et al., 2018, Fries et al., 2007, Miller et al., 2009].

### Effects of stimulation at low and high frequencies

To illustrate the neuronal effects from stimulation at different frequencies, we first show data from an example subject who received electrical stimulation in one location at four frequencies: 10, 50, 100, and 200 Hz. Each frequency was tested 96 times at each amplitude. To measure the effect of stimulation at each frequency on neuronal activity, we computed the mean spectral power in the HFA band at each recording electrode in a 500-ms interval before and after each stimulation trial (Fig. 1A). In many cases we found statistically reliable changes in HFA as a result of stimulation at a particular frequency (e.g., see Figure 1B–C; z = 5.47, p*<*10^−6^, signed-rank test, uncorrected). The HFA changes from stimulation were often present at multiple recording electrodes. This extent of these HFA changes is illustrated in Figure 1D, which shows that this subject had widespread electrodes that showed significantly decreased HFA power when 10-Hz, 1-mA stimulation was applied at a site in the left lateral temporal lobe.

To quantify the changes in HFA power that resulted from each type of stimulation, we computed the mean power change across stimulation trials for each recording electrode (Fig. 1E), excluding sites showing artifacts (see *Methods*). For this site, 10-Hz stimulation at 1 mA caused a significant decrease in mean HFA power across electrodes (z = *−*7.59, p*<*10^-10^, signed-rank test, uncorrected; Fig. 1E). Notably, the recording electrodes that showed significant changes in HFA power included locations both proximal and distal to the stimulation site, even in contralateral areas (Fig. 1D), which might be considered surprising in light of previous studies that focused on the local effects of stimulation [Limousin et al., 1995, Dostrovsky et al., 2000, Logothetis et al., 2010].

We next examined whether a similar pattern of HFA changes was present for stimulation at other frequencies in this subject. Figure 1B shows the pattern of HFA power changes that resulted from 200-Hz, 1-mA stimulation at this same site. In contrast to the 10-Hz stimulation, here we instead found HFA power increases (Fig. 1I). This HFA power increase was robust at the level of individual electrodes (Fig. 1H; z = 5.03, p*<*10^-5^, signed-rank test, uncorrected) as well as at the group level across this subject’s brain (Fig. 1J; z = 4.64, p*<*10^-5^, signed-rank test, uncorrected). Thus, the data from this subject illustrate that the effect of stimulation can be frequency dependent, with 10- and 200-Hz stimulation at the same site and amplitude having opposite effects on HFA power. Because we also found similar patterns of results in other subjects (Fig. S1), we next characterized this effect at the group level.

### Population analysis of the effects of stimulation frequency and amplitude

To characterize the effects of stimulation with different parameters across our dataset, we computed the proportion of all recording electrodes that showed significant HFA decreases or increases for each unique combination of stimulation site, frequency, and amplitude. Figure 2A illustrates, for each stimulation parameter, the percentage of recording electrodes that showed significant HFA power decreases averaged across stimulation sites. HFA decreases were most prevalent for stimulation at low frequencies and high amplitudes. This pattern was present for both depth and surface stimulation sites. When stimulating surface electrodes at high amplitudes, HFA decreases were prevalent for all frequencies.

To assess the reliability of these effects statistically, we used a linear mixed-effects (LME) model to analyze how the prevalence HFA changes depend on the parameters used for stimulation (see *Methods*). Due to our clinical data collection environments, our dataset is heterogeneous, with individual subjects having variable numbers of stimulation sites and individual sites being stimulated at different frequencies and amplitudes. LME modeling is well-suited for analyzing this type of heterogeneous dataset because it can identify linear trends (including interactions) across multiple factors and can accommodate both repeated and missing measurements [Baayen et al., 2008]. We used the LME model to analyze the distributions of HFA power changes across the dataset (Fig. 2A), and the results confirmed that the frequency and amplitude dependence of HFA power decreases mentioned above were statistically reliable for both depth electrodes (all z′*s*=3.39–4.87; all *p*’s*<* 10^−3^ for effects of frequency, amplitude, and their interaction) and surface electrodes (z′*s*=1.9–3.34; all *p*’s*<* 0.05, see Table S4).

We also used the LME model to examine the parameter dependence of stimulation-induced HFA power increases. Figure 2B shows the mean percentages of recording electrodes that showed significant HFA power increases following stimulation at various parameters. Stimulation on depth electrodes at high frequencies and high amplitudes was most closely linked to increases in HFA power. The LME model confirmed that this effect was robust for depth electrodes, by showing significant effects of stimulation frequency on HFA power as well as a frequency *×* amplitude interaction (both *p*’s *<* 0.05, see Table S4). This finding that higher stimulation currents are associated with broader HFA power increases is consistent with the earlier finding that higher currents are associated with more widespread phosphenes in the visual cortex [Winawer and Parvizi, 2016]. In contrast, for surface electrodes, HFA increases were most prevalent for high-frequency stimulation and low amplitudes (all z′*s*= 0.82-1.80; *p′s >* 0.05 see Table S4).

Figure 2C summarizes these results. Overall HFA decreases were more prevalent than increases, regardless of stimulation frequency and electrode type. Further, stimulation on depth electrodes at high and low frequencies, respectively, was associated with HFA increases and decreases (LME model: HFA increase/decrease *×* Frequency: z = 3.55; p = 0.0004). Notably, for stimulation on surface electrodes, we observed different patterns of frequency dependence for high versus low amplitudes. Whereas high-frequency surface stimulation at high amplitudes rarely caused HFA increases, at lower amplitudes, high-frequency stimulation often caused HFA power increases (see above LME model results).

While these trends were robust statistically, we observed that the HFA power changes showed variability across individual stimulation sites (e.g., Fig. 2D). To measure this variability, we quantitatively compared HFA response patterns across different stimulation sites in the same subject. On average, only 16% of subjects showed similar (positively correlated) patterns of HFA power changes in response to different stimulation sites (Fig. S3A), which supports our approach of separately analyzing individual stimulation sites. Nonetheless, to confirm that our results were not affected by treating individual stimulation sites independently, we also performed the above analyses at the level of each subject, by averaging response patterns across the stimulation sites in each subject prior to population-level statistical analysis. This subject-level analysis confirmed our primary results of a frequency-dependence of HFA power changes (Fig. S3B-E). More broadly, the variability between HFA changes caused by different stimulation sites in a subject suggests that it is important to understand the role of location in modulating neuronal activity.

### Distance to white matter mediates the effects of stimulation

Previous studies showed different neurobehavioral changes from applying stimulation in white versus gray matter [Mayberg et al., 2005, Titiz et al., 2017]. Modeling and animal studies demonstrated that bipolar stimulation creates an electrical potential field between and around the anode and cathode of the stimulation site that activates elements within the activated volume [McIntyre et al., 2004b, Histed et al., 2009, Lujan et al., 2013]. Based on these models, we hypothesized that stimulation applied in proximity to white-matter tracts would have different neuronal effects compared to stimulation in gray matter.

To compare the physiology of white-versus gray-matter stimulation on a large scale, we investigated how the proximity of the stimulation site to white matter correlates with the resulting change in HFA power. We first classified each depth stimulation site according to whether it was in white or gray matter, based on its mean proximity to white matter tracts (see *Methods*), and separately compared the HFA changes for each group. Figures 3A and B show data from two patients who were each stimulated at two nearby sites, one labeled as white matter (labeled # 1) and labeled as gray matter (# 2). Both subjects showed HFA decreases when stimulation was applied at the gray-matter site and, inversely, HFA increases for stimulation at the white-matter site.

**Figure 3:**
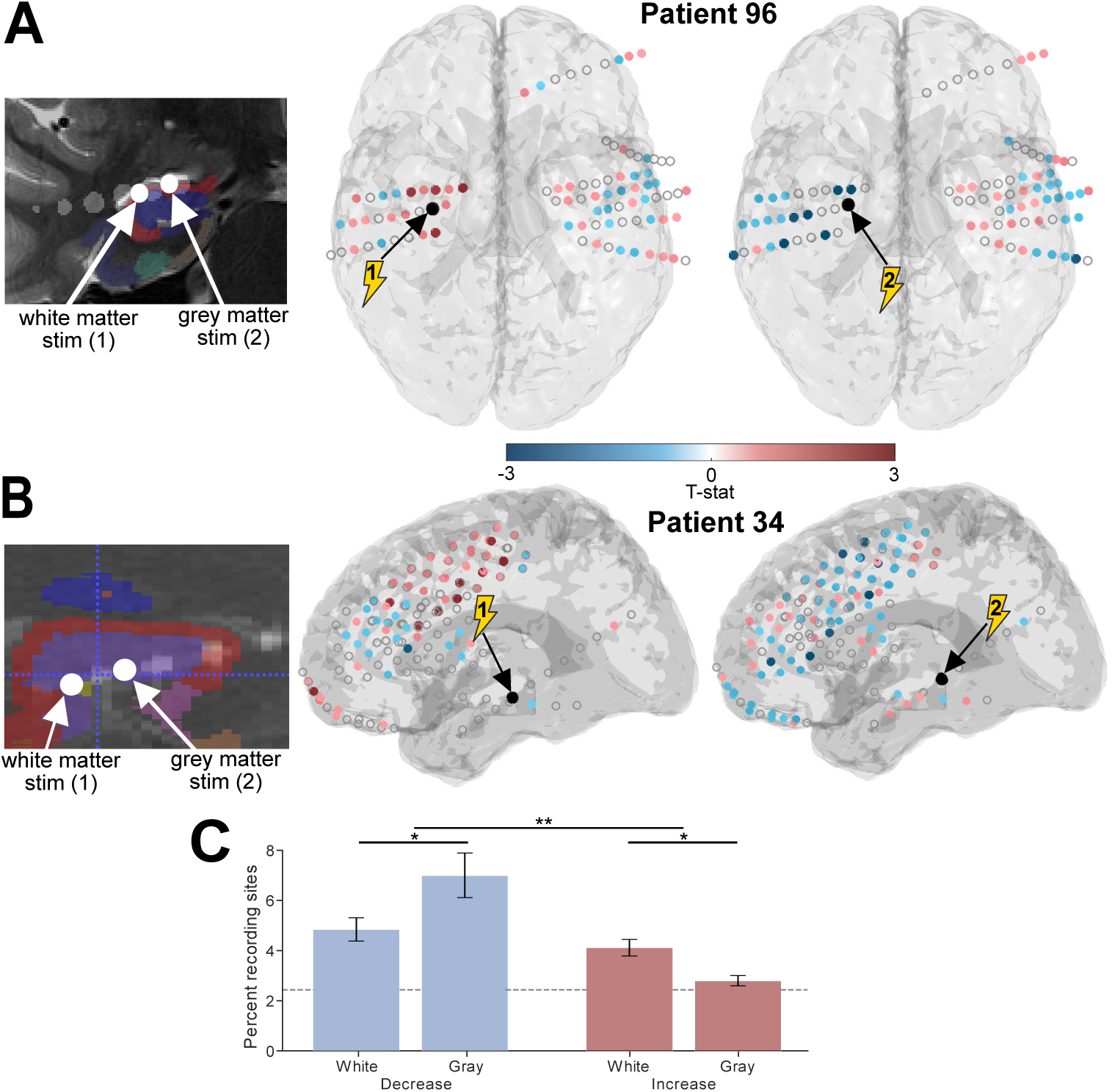
Role of white-matter proximity in modulating the effects of stimulation. **(A)** Brain maps of HFA responses in example Patient 96 of HFA. Stimulation site is indicated in black and color indicates t-statistic of HFA change for recording electrodes. Both sites were stimulated at 200Hz and 0.75mA. Left brain map indicates data for a stimulation site near white matter, which caused significant HFA power increases on 4 recording electrodes (dark red). The right brain map shows data from stimulation at a site in gray matter, which caused a significant decrease in HFA power on 8 recording electrodes (dark blue). Far left panel, coronal MRI image showing the precise location of these two stimulation sites, labeled 1 and 2 corresponding to the left (white) and right (gray) brain maps, respectively. **(B)** Brain map of HFA responses to stimulation near white and in gray matter in example Patient 34. Both sites were stimulated at 200 Hz and 1 mA. Plot format follows panel A. **(C)** Group-level analysis, illustrating the percent of recording electrodes across the entire dataset that showed significant HFA power increases and decreases for white- and gray-matter stimulation. Gray dashed line indicates chance. Error bars: ±1 SEM. *p<0.05, **p<0.01.

We next performed a group-level analysis of the relation of white and gray matter on HFA changes from stimulation. We focused this analysis on stimulation parameters in the range of 100–200 Hz and 0.5–1 mA, which were chosen as the parameters most likely to cause HFA increases. We then compared the prevalence of HFA power changes across sites in white (n=70) and gray matter (n=61). Stimulation at white-matter sites caused a greater rate of HFA increases compared to sites in grey matter (Fig. 3C). Inversely, gray-matter stimulation caused HFA power decreases at more sites compared to white-matter stimulation. Analyzing the prevalence of each type of HFA change with a two-way ANOVA, we confirmed that there was a statistically significant interaction between the white- or gray-matter location of stimulation and the prevalence of HFA increases and decreases (F(1,1) = 6.55; p = 0.01).

### Spatial spread of neuronal activity changes from stimulation

We next examined the spatial spread of stimulation-induced changes in HFA. To do this, we measured the prevalence of HFA increases and decreases as a function of recording electrodes’ distance from the stimulation site. Overall, the prevalence of HFA power decreases was greater for recording electrodes near the stimulation site compared to distal electrodes (Fig. 4A–D; Fig. S1). A similar distance dependence was present for recording electrodes that showed HFA increases. Although HFA increases were generally less prevalent than decreases, the prevalence of HFA decreases fell off more drastically with distance to the stimulation site as compared to HFA increases (LME model: Distance *×* Direction interaction: *z* = 5.62, *p <* 10^−9^, see Table S4).

**Figure 4:**
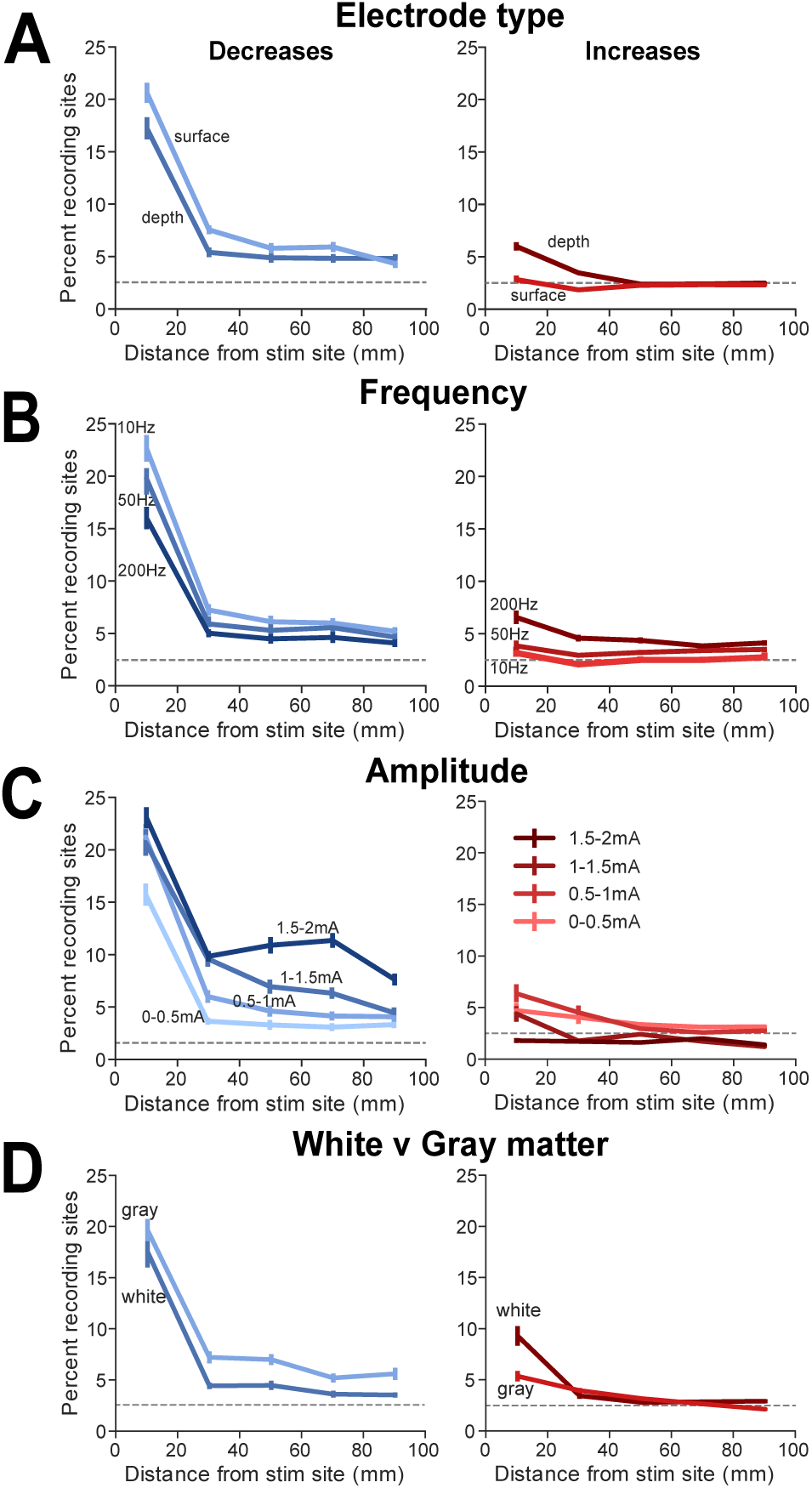
Spatial spread of neuronal activity changes. All plots show the mean percent of recording electrodes that showed significant HFA power decreases (left) and increases (right) binned by their distance from the stimulation site. **(A)** Comparison of effects between depth and surface stimulation sites. Gray dashed line indicates chance. All error bars denote: ±1 SEM. **(B)** Analysis for effects of stimulation frequency (10, 50, and 200 Hz.) **(C)** Analysis for effects of stimulation amplitude (0– 0.5, 0.5–1, 1–1.5, & 1.5–2 mA). **(D)** Analysis for effects of stimulation near white versus gray matter (see Methods).

We compared the spatial spread of HFA increases and decreases separately for depth and surface stimulation (Fig. 4A). Stimulation at both depth and surface sites showed that the prevalence of HFA decreases diminished with distance at approximately the same rate, but HFA decreases from surface stimulation are more prevalent across the brain (Depth vs. surface: F(1) = 5.52, p=0.01; Distance *×* depth/surface interaction: F(4,1) = 1.21, p=0.30, two-way ANOVA). Inversely, HFA power increases from depth stimulation were more prevalent and showed a distance effect than increases from surface stimulation (Depth vs. surface: F(1) = 7.77, p=0.005; Distance *×* depth/surface interaction: F(4,1) = 2.25, p=0.06, two-way ANOVA).

Next we examined the role of stimulation frequency on the distance dependence of HFA power changes (Fig. 4B). For all frequencies, HFA power decreases were most prevalent at recording electrodes near the stimulation site. This effect was significantly larger for stimulation at low frequencies (LME model: Distance *×* Frequency: *z* = *−*4.26, *p* = 0.00002). A related drop-off with distance was also present for the sites that showed HFA power increases (right panel); however, this effect was most prevalent for 200-Hz stimulation (Distance *×* Frequency: *z* = *−*2.72, p = 0.006, LME model).

We also examined the role of stimulation amplitude in the distance dependence of HFA changes (Fig. 4C). As in the above analyses, the prevalence of HFA changes decreased with distance from the stimulation site. However, the rate of this fall-off inversely correlated with stimulation amplitude. For low stimulation amplitudes, HFA decreases were present at *∼*5% electrodes with distances *≥*30 mm from the stimulation site, but for amplitudes at *≥*1 mA, *∼*10% of electrodes spaced at *≥*30 mm showed HFA decreases. The interaction between distance and amplitude had a statistically significant effect on the prevalence of HFA decreases (Distance *×* Amplitude interaction: z = −3.08; p = 0.002, LME model). This indicates that larger stimulation amplitudes increase the spatial spread of stimulation-induced HFA decreases. This type of distance dependence was not evident in the sites that showed HFA increases from stimulation (Fig. 4C, right panel).

Finally, we analyzed the spatial spread of HFA power changes from white-versus gray-matter stimulation (Fig. 4D). This analysis showed that the spatial spread of HFA decreases was more prevalent across the brain when stimulation was applied near gray matter matter (left panel: White vs. Grey Matter: F(1) = 4.46, p = 0.04; Distance *×* Matter interaction: F(4,1) = 0.41, p=0.8, two-way ANOVA), and an opposite effect was present for HFA increases, which were more prevalent when stimulating near white matter (right panel: White vs. Grey Matter: F(1) = 4.63, p = 0.03; Distance *×* Matter interaction: F(4,1) = 1.15, p=0.33, two-way ANOVA). We have confidence that our results are not due to limitations of the recording system because the presence of our effects differ with the precise location of the electrodes with regard to the brain, which is unlikely to be related to the electrical characteristics of the recording and stimulation systems. In particular, given HFA increases are greater for stimulation on depth electrodes near white matter than other areas, it means that the effect is likely related to physiological differences rather than stimulation artifact. More broadly, because our results show different effects for white-versus gray-matter stimulation, it suggests that clinicians should select stimulation sites based on tractography to bring about desired changes in neuronal activity.

### Stimulation-induced resetting of neuronal activity

In addition to identifying HFA power increases or decreases from stimulation, we also observed a different phenomenon, in which stimulation caused HFA power to adjust to a fixed level. In contrast to the above-described sites that showed increases or decreases in mean HFA power after stimulation, an electrode that exhibited “HFA resetting” would show variable HFA power prior to stimulation across trials that became tightly clustered after stimulation. Therefore, to identify this phenomenon we compared the variances of HFA power at each electrode between pre- and post-stimulation intervals (rather than comparing the means as in earlier analyses). Figure 5A shows two example left temporal-lobe recording electrodes that exhibited resets in HFA power from stimulation. Each of these electrodes showed substantial variation in HFA power before stimulation, with this variation decreasing significantly (both *p*’s*<* 10^−6^) afterward (Fig. 5B). The data in this figure illustrate two characteristics of resetting: First, that the recording electrodes that show HFA power resets are often spatially clustered. Second, that HFA resetting is not necessarily found immediately surrounding the stimulation site, which could have been indicative of artifact.

**Figure 5:**
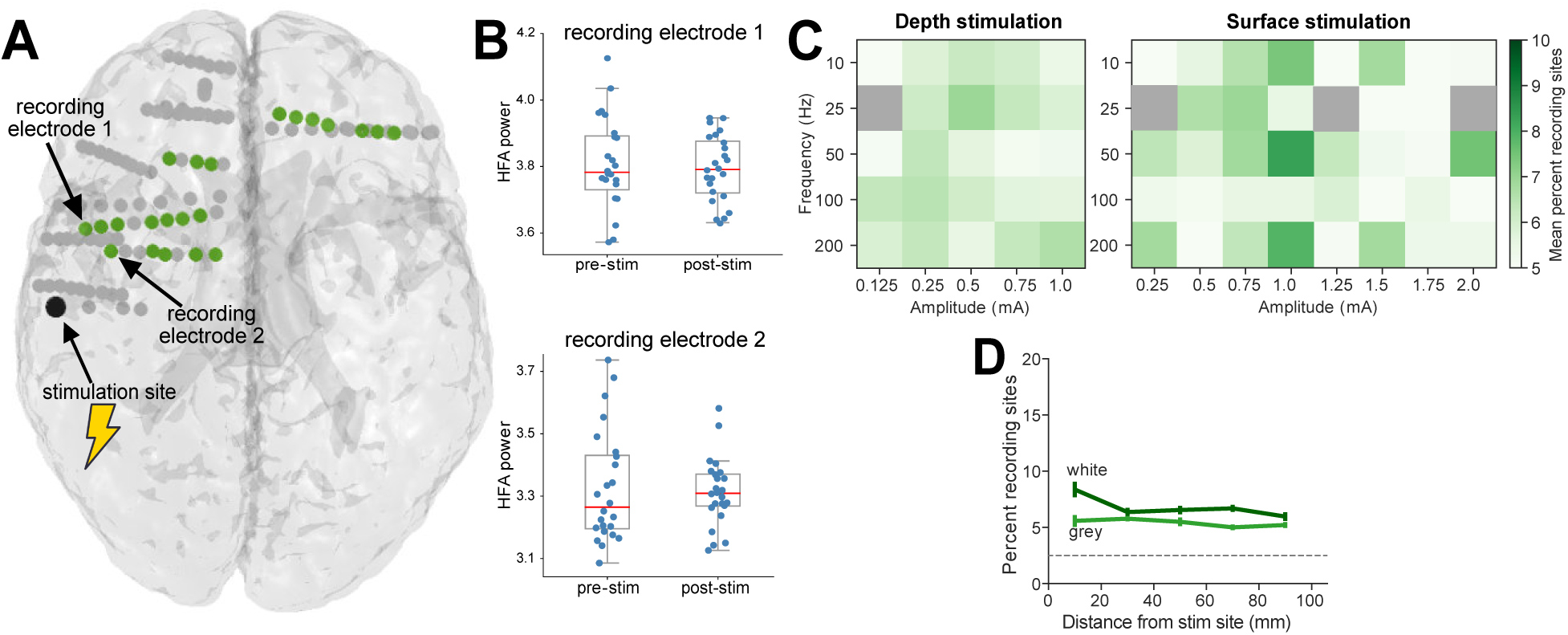
Stimulation induced HFA power resetting. **(A)** Brain map from Patient 200 illustrating the recording electrodes that showed significant power resetting (green) following stimulation at one labeled site (black). **(B)** HFA power measured from two example recording electrodes in this patient before and after stimulation. Both sites show significant resetting, in which the variance of HFA power significantly decreases from pre-to post-stimulation without a significant change in the mean power (contact 1: p<10^-6^, uncorrected F - test; contact 2: p<10^-8^). **(C)** Group-level analysis showing the overall mean percent of recording electrodes that showed significant HFA resetting for each combination of parameters. **(D)** Mean percent of recording electrodes that showed significant power resetting as a function of distance from white- and gray-matter stimulation sites.

To statistically characterize HFA resetting, we identified the recording electrodes that showed a significant decrease in the variance of HFA power from pre- to post-stimulation (*F* test, *p <* 0.05) with no change in mean (*t* test, *p >* 0.05). Analogous to the above analyses, we computed the proportions of electrodes that showed significant resetting for each combination of stimulation frequency and amplitude (Fig. 5C). This analysis suggested that HFA resetting for each stimulation site is dependent on stimulation frequency (both p,*s <*10^-4^, for depth and surface stimulation, see Table S4). The LME model did not show a significant dependence for the prevalence of HFA resetting according to the stimulation amplitude (Depth: p = 0.22; Surface: p = 0.61) or the interaction between frequency and amplitude (Depth: p = 0.24; Surface: p = 0.08).

We also examined the prevalence of HFA resetting as a function of distance to the stimulation site. HFA resetting was greater at recording electrodes closer to the stimulation site. For electrodes near the stimulation site, the prevalence of resetting was significantly less than that of HFA power decreases and greater than the prevalence of HFA power increases (LME model, Distance *×* Resetting vs. Increase vs. Decrease: z = 2.4, p = 0.007; Fig. 5D). Additionally, we found that the prevalence of HFA resetting was greater for stimulation in white rather than gray matter (White vs. Grey Matter: F(1) = 4.01, p = 0.04; Distance *×* Matter interaction: F(4,1) = 0.58, p=0.67, two-way ANOVA). In light of its distinctive characteristics, these results indicate that stimulation-induced HFA resetting reflects a distinctive neuronal phenomenon compared to stimulation-induced HFA power increases and decreases.

### Control analyses of stimulation artifact effects

While one cannot completely separate artifact from physiological signals in clinical iEEG recordings, we took a two-stage approach to identify and mitigate their potential impact on our results. As described in the *Methods*, we ensured that electrical artifacts from the activation of the stimulator did not impact our HFA power calculations by measuring HFA using temporally precise multitapers at an interval that was separated in time from when the stimulator was active. As shown in Figure S4, this approach successfully identified reliable patterns of HFA power increases that had different timecourses compared to stimulation artifacts.

We also examined whether our results were affected by artifacts related to amplifier saturation. After stimulation concludes, on many electrodes the iEEG recording shows a transient low-frequency deflection. This type of deflection could disrupt accurate power measurement. To minimize the influence of this type of artifact on our results, as described in the Methods we removed both individual trials and recording electrodes that exhibited large post-stimulation voltage changes (Fig. S2). To further validate that our results were not correlated with this kind of artifact, we also performed the above population analysis (Fig. 2A,B) using three different artifact-rejection thresholds (Fig. S6). The relationship between HFA changes and stimulation parameters remained present for all thresholds (Table S4). This indicates that the HFA changes we found are not a result of post-stimulation artifacts because these artifacts were removed at different rates across thresholds. We also measured the prevalence of artifacts for each combination of stimulation amplitude and frequency (Table. S3). Because artifact rates, unlike HFA, did not substantially vary across stimulation parameters, it additionally supports our view that the frequency dependence of HFA changes we observed was not a result of stimulation artifacts.

## Discussion

Clinicians and researchers are increasingly interested in brain stimulation because it provides a way to directly modulate ongoing brain activity to support various goals including treatment of neurological disorders. However, for brain stimulation to be used optimally, stimulation should be targeted precisely according to the desired outcome. One goal of our project was to guide selection of stimulation parameters by characterizing—across space, frequency, and amplitude—the neuronal effects of direct cortical stimulation in humans. Our work indicates that effects of stimulation significantly differ depending on the parameters used for stimulation. There were also substantial variations in the effects of stimulation across subjects. Together, our results indicate that we may achieve more effective outcomes for stimulation by choosing parameters according to the desired neuronal pattern.

A key result from our work is demonstrating that the neuronal effects of direct brain stimulation in humans are frequency dependent. While the general effect of stimulation on HFA was negative, we demonstrated that high- and low-frequency stimulation inversely impact neuronal activity, preferentially causing HFA power increases and decreases, respectively (Fig. 2C). In this way, our work extends prior studies that demonstrated that the frequency of stimulation was an important factor in driving specific clinical outcomes from stimulation. For example, when using DBS for Parkinson’s disease, stimulation at frequencies over 90 Hz alleviated tremor while frequencies below 60 Hz aggravated tremor [Ushe et al., 2004, Fogelson et al., 2005, Kuncel et al., 2006]. Further, there is evidence of frequency dependence in the use of stimulation to treat epilepsy, in which stimulating at frequencies below 2 Hz and above 70 Hz reduced epileptic activity, whereas intermediate frequencies had no effect [Mina et al., 2013, Yu et al., 2018]. With these findings and others, our work indicates that stimulation frequency should be tailored according to the goals of the procedure.

The frequency dependence we observed is generally consistent with findings from animals. Of particular relevance to our work is the study by Logothetis et al. [2010] who measured the resultant changes in neuronal activity in various brain regions of monkeys following microstimulation at a range of frequencies. Consistent with our results, that study found that low-frequency stimulation caused decreases in neuronal activity whereas high-frequency stimulation caused mixed increases and decreases in different downstream regions. It is notable that the findings from that study converged with ours despite the substantial methodological differences. Whereas we applied stimulation at macroelectrodes in human patients and measured HFA power, the Logothetis et al. [2010] study used microstimulation in normal monkeys and measured fMRI and single-neuron spiking.

A question that arises from these results is why stimulation at low frequencies suppresses and stimulation at high frequencies is more likely to activate. Quantitative models suggest that high-frequency stimulation selectively activates fibers of passage and axon terminals with low thresholds that would not normally be activated by low-frequency stimulation [McIntyre and Grill, 2002]. This may occur because high-frequency stimulation delivers a higher rate of charge with shorter time between pulses, which increases mean spiking rates because neurons have less time to hyperpolarize [Ranck Jr, 1975, Benazzouz and Hallett, 2000, Jensen and De Meyts, 2009, McIntyre et al., 2004a]. By additionally incorporating neuroanatomy, models may also explain our finding of prevalent HFA decreases near the stimulation site, while HFA increases were relatively more widespread (Fig. 4A–C). These spatial variations may be explained by the anatomical organization of the stimulated neurons. When stimulation activates axons, which is more likely with high frequencies [McIntyre and Grill, 2002]; models suggest that the excitatory effects can spread more broadly, following axonal projections to other regions. Inversely, when stimulation impacts cell bodies, the effects are likely to be inhibitory and spatially limited [McIntyre and Grill, 2002, McIntyre et al., 2004a, Herrington et al., 2015].

It is notable that we found variability in HFA power changes between stimulation sites even within an individual. This result is consistent with the idea that local and distal effects of stimulation depend on the neuronal morphology surrounding the stimulation site [Pouratian et al., 2004, Lesser et al., 2008, Borchers et al., 2012]. For instance, the effects of stimulation may depend on the precise positioning of the implanted electrode and its specific orientation relative to cortical layers or fibers of passage. At the broadest level, our findings support the idea that the effective use of brain stimulation should consider neuron organization, thresholds, and neurotransmitters of an area to better predict the downstream effects of stimulation [Ranck Jr, 1975]. This variation that we found in the responses to stimulation at different sites might help explain prior studies that showed diverse perceptual and behavioral responses to stimulation between subjects and stimulation locations [Selimbeyoglu and Parvizi, 2010, Borchers et al., 2012, Pouratian et al., 2004]. Despite this variability, in 16% subjects, we found significantly correlated patterns of HFA power changes across different stimulation sites. This suggests that some individuals have distinctive neuroanatomical patterns, perhaps involving connectivity or genetic differences [Fox et al., 2005], that causes them to show consistent HFA changes even across widespread distributed stimulation targets.

We found that inhibitory and excitatory effects were relatively more likely from stimulation in gray and white matter, respectively. This result adds to a growing body of literature emphasizing that behavioral and electrophysiological outcomes depend on the proximity of stimulation to structural connections. In particular, many studies showed that positive behavioral outcomes result from stimulation in white rather than gray matter. In particular, studies reported improvement of memory specificity and depression symptoms when applying stimulation to white matter rather than gray matter [Suthana et al., 2012, Titiz et al., 2017, Mayberg et al., 2005, Gutman et al., 2009]. Similarly, one recent study showed that white-matter stimulation amplifies oscillatory theta coherence across memory networks [Solomon et al., 2018]. Additionally, studies in rodents show similar results, demonstrating that microstimulation in white matter was more effective for exciting distal neuronal populations [Nowak and Bullier, 1998a,b]. Our findings add to this body of work, by suggesting a mechanism for white-matter stimulation to improve behavior, by preferentially causing neuronal excitation. Recent modeling studies determined patient-specific stimulation locations based on predictions of electrical-field generation based on patient tractography [Lujan et al., 2013]. Going forward, it may be beneficial for clinicians to integrate patient-specific models to guide stimulation locations relative to relevant structural connections.

Besides using stimulation to excite and inhibit, we observed the novel phenomenon of stimulation-induced HFA resetting. Some prior closed-loop stimulation studies continuously monitored the current brain state and delivered stimulation to increase or decrease a particular measure of neuronal activity when it crossed a critical threshold [Ezzyat et al., 2017, Sun and Morrell, 2014]. In contrast to this approach of using stimulation to shift neuronal activity in one direction, the existence of stimulation-induced resetting indicates that targeted stimulation can induce a specific state regardless of the level of neuronal activity prior to stimulation. By leveraging stimulation-induced resetting, we hypothesize that targeted white-matter stimulation protocols can transition brain activity into particular states [Stiso et al., 2018], supplementing existing closed-loop methods that focus on shifting ongoing neuronal patterns in one direction.

Although we conducted our work with electrodes implanted in surgical patients, our results also have implications for non-invasive brain stimulation. Much like direct electrical stimulation, transcranial magnetic stimulation (TMS) and transcranial electrical stimulation (TES) have been shown to produce mixed excitatory and inhibitory responses. The direction of the changes in neuronal activity caused by TMS and TES were shown to depend on parameters that were analogous to those we tested, such as the location, frequency, and amplitude of stimulation [Hallett, 2000, 2007, Barker and Shields, 2017, Fertonani and Miniussi, 2017, Antal and Paulus, 2013]. Furthermore, non-invasive brain stimulation studies also found substantial inter-subject variability [Ĺopez-Alonso et al., 2014, Wiethoff et al., 2014], which is also consistent with our results. Given these similarities, our results support the approach of customizing non-invasive stimulation parameters for each individual, as we found with invasive stimulation.

Although electrical stimulation can cause substantial artifacts in neural recordings, we have reason to believe that artifacts are not driving our results. We applied an established method of artifact rejection (see *Methods*; Solomon et al. [2018]) and showed that our main results persist irrespective of the particular level of artifact rejection that we applied (Fig. S6). An additional reason we have confidence that our results reflect neural signals is because our characterization of HFA changes matches the frequency dependence seen in animals [Logothetis et al., 2010]. Additionally, the stimulation-induced HFA changes we found also interact with neuroanatomy—HFA increases were more prevalent when stimulating white rather than gray matter—which is a pattern that is unlikely to appear as the result of electrical artifacts.

A focus of many types of brain stimulation is to recapitulate a target neuronal pattern. Because we show the stimulation parameters that cause different types electrophysiological signals, our work provides a guide for clinicians to help select stimulation frequencies and amplitudes that recreate particular target patterns. In this regard, the most important features of our results are (1) that high- and low-frequency stimulation are associated with HFA power increases and decreases, respectively, and (2) that high stimulation currents cause HFA power decreases across broader cortical regions. These patterns also help explain key features of previous neuromodulation work. For example, in one study we found that stimulation at a particular site caused a patient to spontaneously recall an old autobiographical memory, and, notably, this site showed HFA decreases when that memory was remembered normally [Jacobs et al., 2012]. Our current findings help explain why this occurred, because the 50-Hz stimulation that was used was likely to cause an HFA power decrease that matched the neuronal pattern associated with that memory. Further, our results help explain the recent finding that high-frequency stimulation in the lateral temporal lobe can help improve episodic memory encoding [Kucewicz et al., 2017, Ezzyat et al., 2017]. Normally, successful learning of episodic memories is associated with elevated HFA power [Burke et al., 2013]. Therefore, our results help explain that high-frequency stimulation improved memory encoding is because it recreated the elevated HFA power that was normally associated with successful encoding. Our findings do not eliminate the current clinical standard of iteratively testing parameters to optimally select patient-specific stimulation parameters. They do, however, provide clinicians with a better starting point for selecting patient-specific stimulation frequencies to evoke a specific neuronal responses.

There is widespread and growing interest in using brain stimulation for various research, clinical, and practical purposes [Borchers et al., 2012, Ezzyat and Rizzuto, 2018]. In many cases, the stimulation parameters that are chosen for a given task are modeled after the ones used in other protocols or in other subjects [Lozano et al., 2019]. Our work supports a tailored approach to choosing stimulation parameters, by customizing the parameters for each person based on how different types of stimulation affects their own ongoing brain signals as well as the electrophysiological pattern of interest. By combining our observations of electrophysiological effects of stimulation with modeling and knowledge of neuronal patterns, clinicians and researchers can design more targeted therapeutic stimulation protocols to more effectively treat neurological and psychiatric disorders.

## Supporting information

Supplemental Analyses

## Acknowledgements

We are indebted to all patients who volunteered their time to participate in our study. We thank Shachar Maidenbaum and Salman Qasim for providing helpful feedback on the manuscript. We thank Blackrock Microsystems and Medtronic, Inc. for providing neural recording and stimulation equipment.

## Author contributions statement

U.M. and J.J. designed and implemented the data analyses and wrote the manuscript. A.W. and J.M advised analysis framework. M.K., and D.R. designed the stimulation-mapping protocol, B.L., M.R.S., G.W., R.E.G, K.Z., B.J., K.D., S.S. recruited subjects, collected data, and performed clinical duties associated with data collection including neurosurgical procedures or patient monitoring. J.S., R.G., and S.D. performed anatomical localization of electrodes. All authors participated in editing.

---

**Figure S1:**
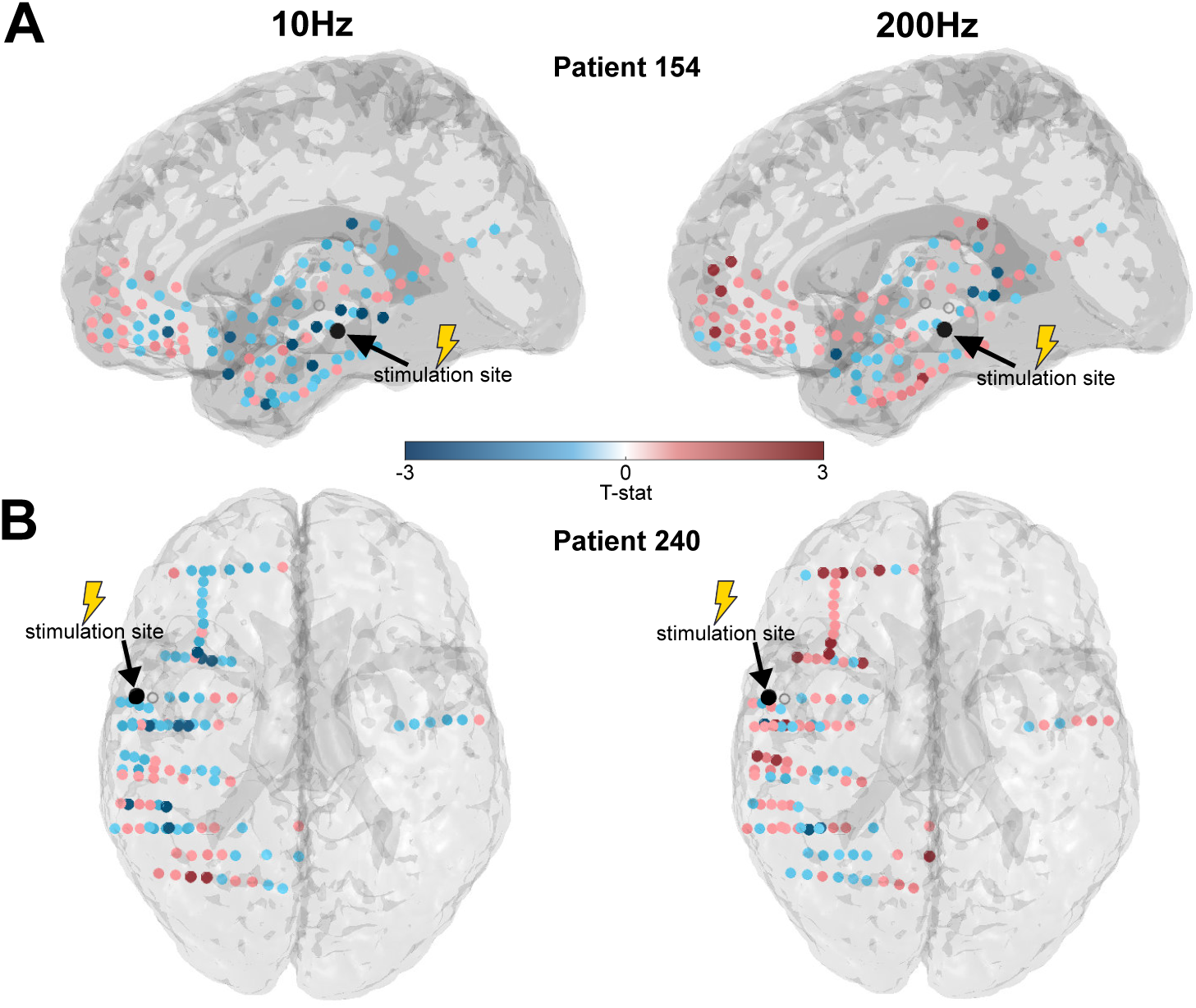
HFA responses depend on stimulation frequency: Additional example subjects. **(A)** Brain maps showing the mean HFA responses across recording electrodes to 10-Hz (left) and 200-Hz (right) stimulation at the same site in Patient 154. The stimulation site is indicated in black and color indicates the t statistic of the change in HFA power from stimulation at each recording electrode. The recording electrodes that are excluded due to artifacts are indicated with an open gray circle. **(B)** Brain maps of HFA responses to 10-Hz (left) and 200-Hz (right) stimulation at the same site in example Patient 240. Plot format follows panel A.

**Figure S2:**
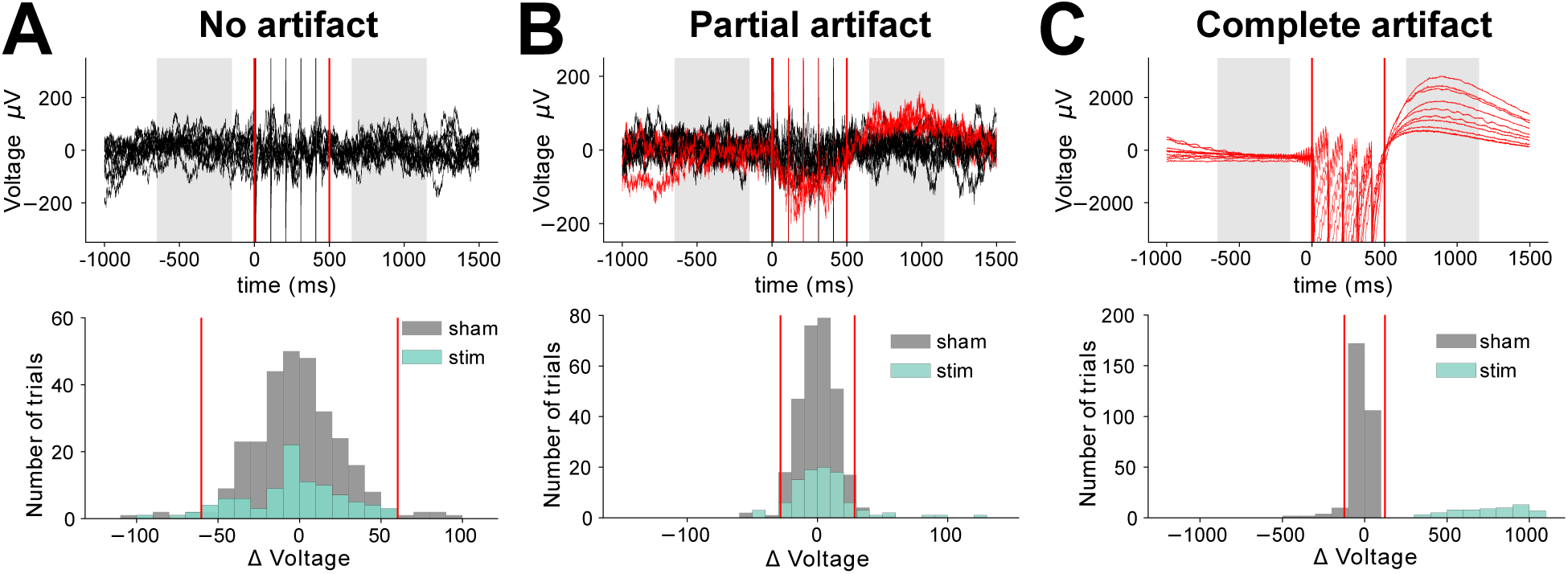
Illustration of our methods for detecting post-stimulation artifacts. **(A)** Top panel, raw signals from 10 trials recorded on an example electrode (Patient 195, electrode 21). Shading indicates the 500-ms time periods before and after each stimulation trial when we measured HFA power. Red lines denote stimulation onset and offset. Bottom panel, histogram of the differences in voltage (post-minus pre-stimulation) for trials when stimulation was applied (turquoise) as well as sham trials (gray). Red lines indicate the artifact-rejection threshold (2SD of sham distribution). Because all of the voltage values on stimulation trials fell within the inner bounds of the thresholds, no trials were rejected. **(B)** An analogous plot to Panel A, created from data of 10 trials on a different example recording electrode (Patient 195, electrode 22). Here, two trials (shown in red) were identified as showing post-stimulation artifacts because their change in voltage (POST-PRE) fell outside the 2-SD threshold computed from the voltage difference measured on that electrode for sham trials (see bottom panel). **(C)** This plot shows data from a third electrode (Patient 195, electrode 18), where all ten trials were identified as showing post-stimulation artifacts. This entire electrode was excluded from subsequent analyses because all trials showed artifacts (see Methods).

**Figure S3:**
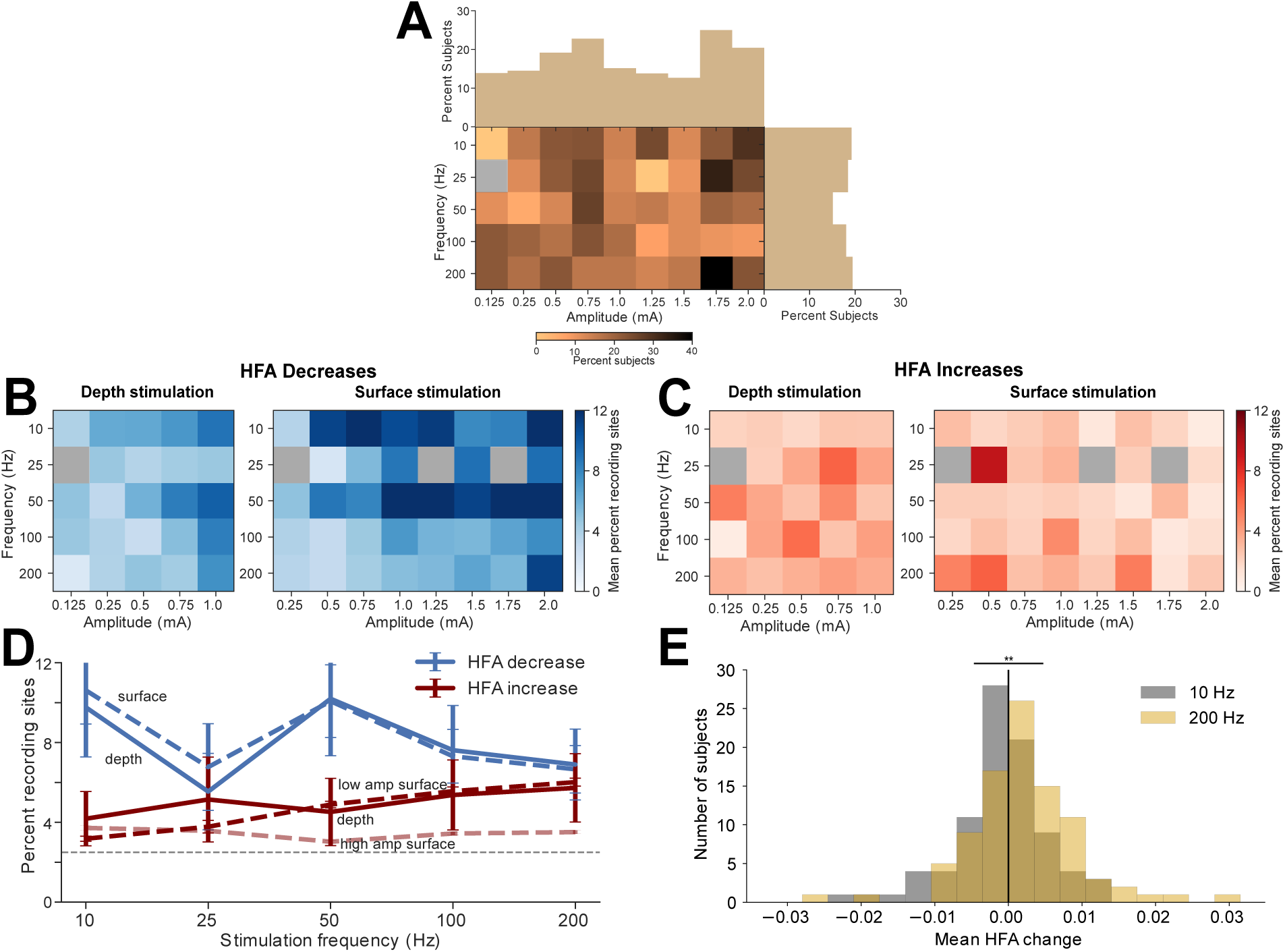
Subject-level analyses of the effect of stimulation frequency and amplitude on HFA power. **(A)** To test whether a subject showed the same response pattern across different stimulation sites, we computed the intraclass correlation coefficent (ICC) between HFA patterns produced by different stimulation sites. A significant ICC indicates that a similar brain-wide HFA pattern was created by stimulating at different locations in the same subject. This plot illustrates, for each frequency and amplitude, the percentage of subjects that showed similar response patterns across different stimulation sites (as identified with a significant positive ICC). This analysis showed that, on average, 16% of subjects show similar HFA patterns for multiple stimulation sites. Because of this above-chance similarity across stimulation sites, we conducted subject-level analyses of the effects of stimulation, rather than stimulation site-level analyses as in Figure 2. **(B)** Subject-level analysis of the mean percent of recording electrodes that showed significant HFA decreases for each combination of stimulation frequency and amplitude, separately computed for depth (left) and surface (right) stimulation. LME modeling shows a similar pattern of statistical effects as in Figure 2A (see Table S4). **(C)** Subject-level analysis of the mean percent of recording electrodes that show significant HFA increases for each combination of stimulation parameters. Again, LME modeling shows similar results as Figure 2B. **(D)** Subject-level analysis that is analogous to Figure 2C. Direction of HFA change × Frequency interaction: z = 3.21; p = 0.0006, LME model). **(E)** Subject-level analysis that is analogous to Figure 2D. These distributions differ significantly (z = *−*3.82, *p <* 10^−3^, *rank-sum test)*.

**Figure S4:**
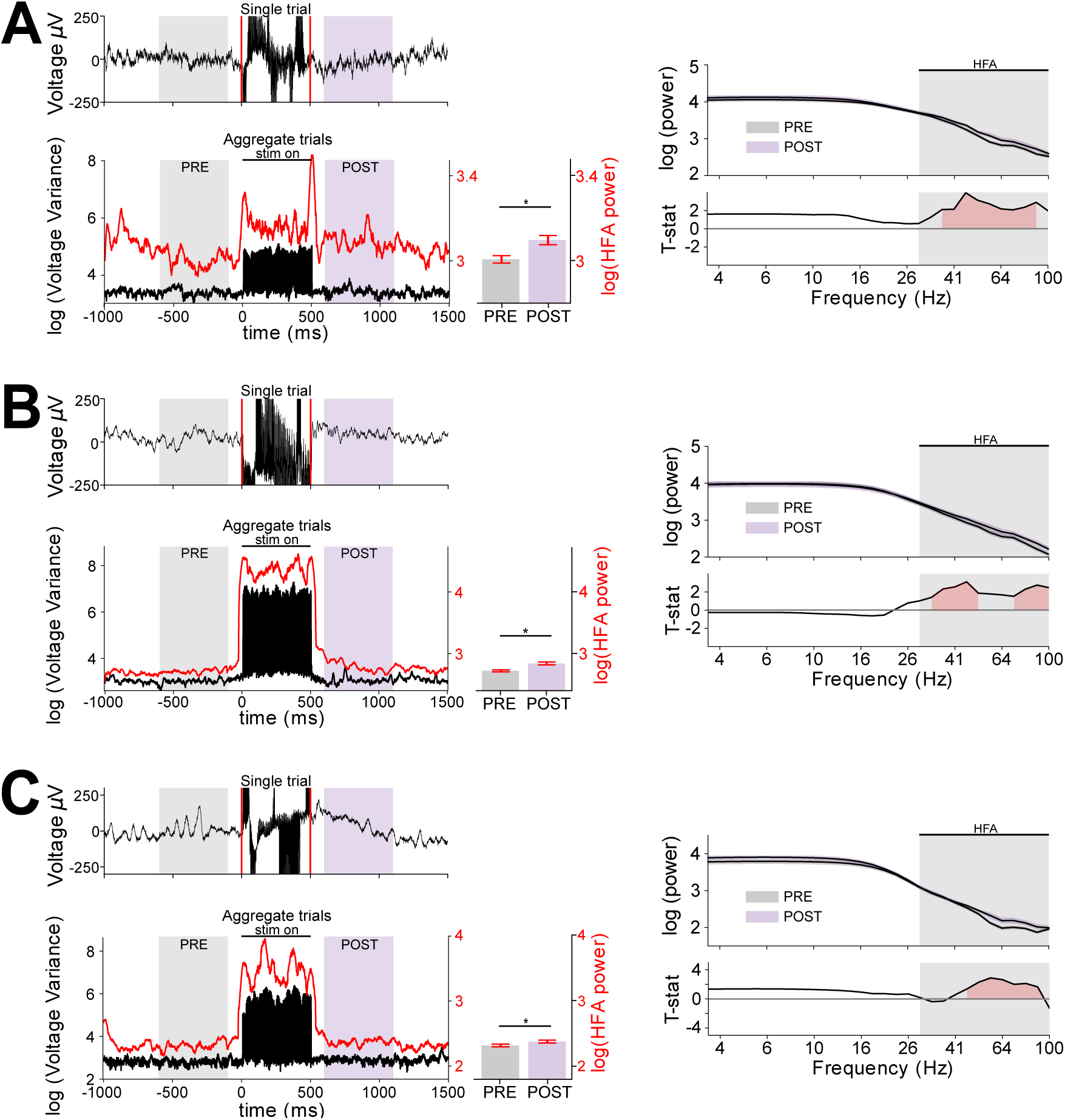
Illustration of how our analysis method avoids high-frequency stimulation artifacts. **(A)** Top-left panel, raw signals (black line) from one trial recorded from Patient 195, Electrode 67. Red lines denote stimulation onset and offset. Bottom-left panel, illustration of the timecourse of stimulation artifacts seen on this channel. Black line indicates the variance of voltage measurements across trials at each timepoint. The marked increase in variance indicates that stimulation artifact affects recordings specifically when the stimulator was active (“stim on”). The red line indicates the mean HFA power. The gray and purple shading indicates the pre and post-stimulation analysis periods. Critically, as this plot shows, the impact of stimulation artifacts on HFA power consistently drops off before and after stimulation and does not overlap with the pre-(gray) or post-stimulation (purple) analysis periods. Right-top panel, log-transformed mean power spectrum for the pre- and post-stimulation intervals. This plot illustrates that stimulation most strongly increases activity in the HFA band (gray shading from 30-100 Hz). Right-bottom panel, t statistic of the difference in pre- and post-stimulation power at each frequency. Red shading indicates positive significant differences at p<0.05. **(B)** Plots follow panel A for Patient 154, Electrode 37 **(A)** Plots follow panel A for Patient 240, Electrode 36

**Figure S5:**
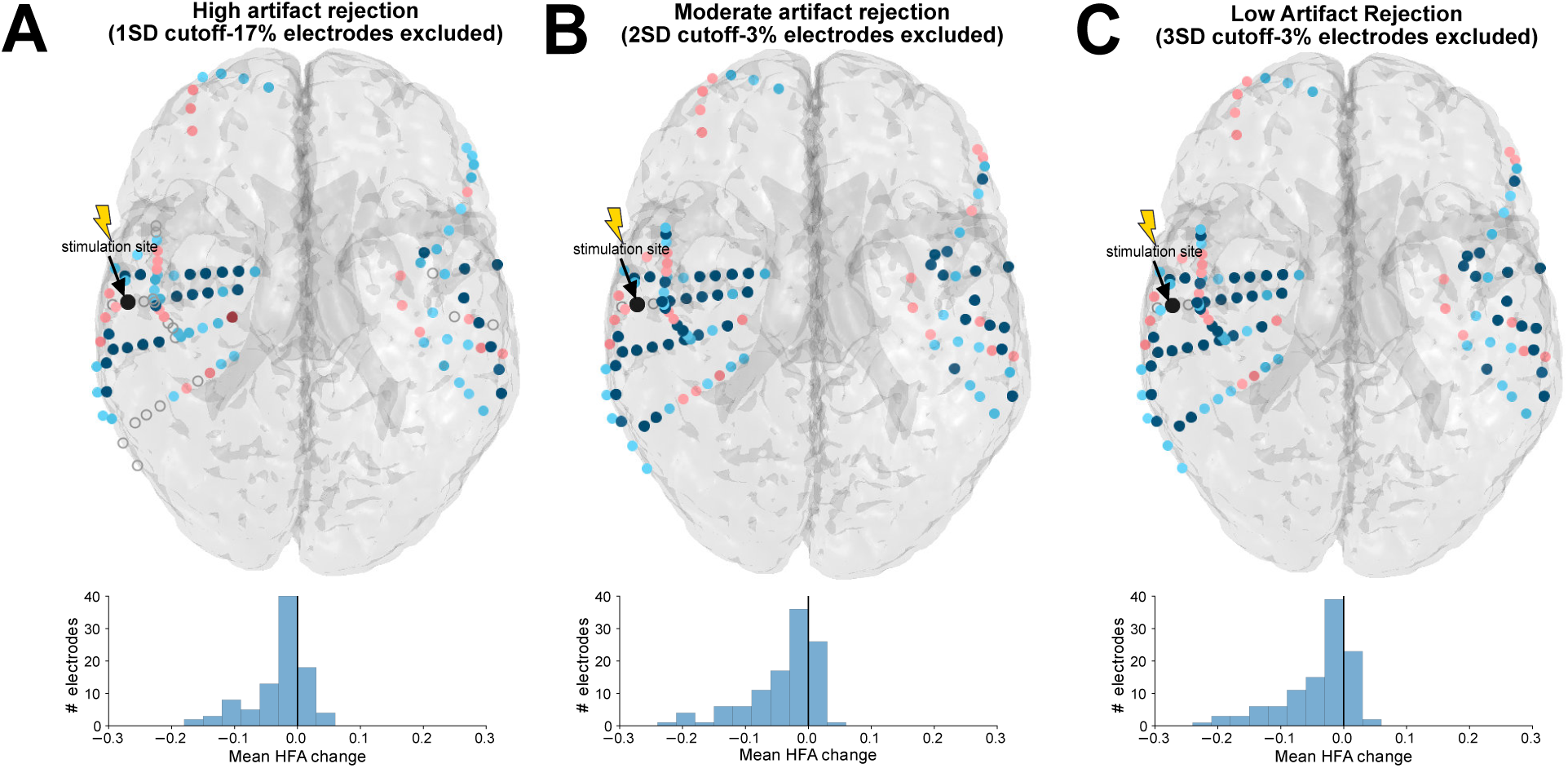
Effects of different artifact-rejection thresholds on HFA power: Data from one example subject. **(A)** Illustration of HFA power changes in patient 195 from 10-Hz, 1-mA stimulation after artifact rejection with a 1-SD cutoff, which excludes 31% of trials and 17% of recording electrodes. Top panel indicates the brain-wide mapping of HFA power changes. Recording electrodes excluded due to artifact indicated are indicated by an open gray circle. Bottom panel, the distribution of mean HFA changes across all analyzed recording electrodes in this subject. **(B)** Analysis of HFA power changes in this patient with a 2-SD cutoff, which excludes 11% of trials and 3% of recording electrodes. **(C)** Analysis of HFA power changes in this patient with a 3-SD cutoff, which excludes 5% of trials and 3% of recording electrodes.

**Figure S6:**
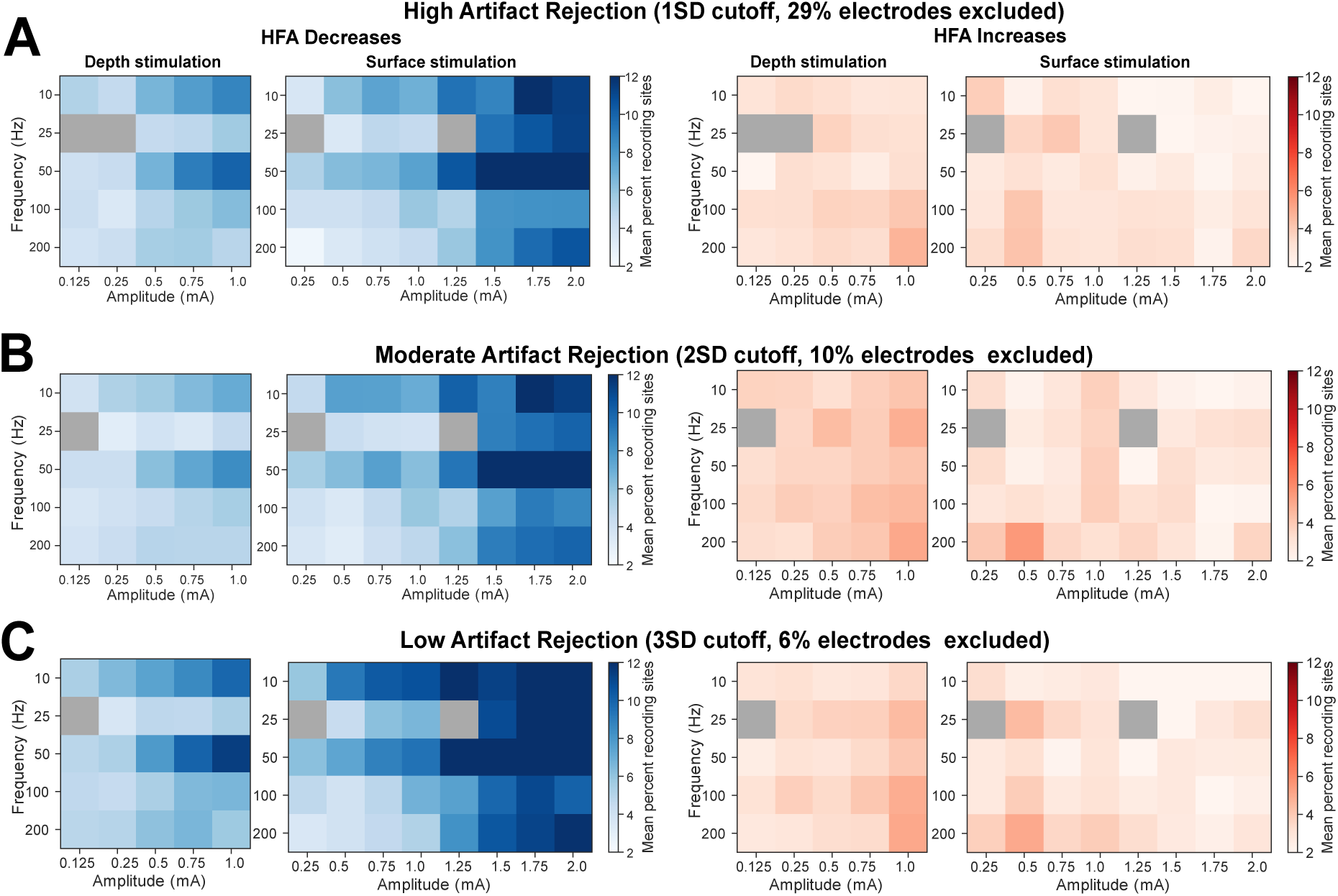
Effects of different artifact-rejection thresholds on HFA power: Population-level analysis. **(A)** Analysis of the percent of recording sites where HFA significantly increased or decreased for when using a 1-SD artifact-rejection threshold. With this threshold we excluded 29% of recording electrodes and 28% of stimulation trials on the remaining electrodes. LME model analysis confirmed a similar relationship between HFA changes and stimulation parameters as in Figure 2A,B (see Table S4). **(B)** Same analysis as our main population results in Figure 2 using a 2-SD artifact-rejection threshold. With this threshold, we excluded 10% of recording electrodes and 12% of trials on the remaining electrodes. **(C)** Same analysis as above, but using a 3-SD artifact-rejection threshold. Here, we excluded 6% of recording electrodes and 5% of trials on remaining electrodes. LME model analysis confirmed a similar relationship between HFA changes and stimulation parameters as in Figure 2A,B (see Table S4).

**Table S1:**
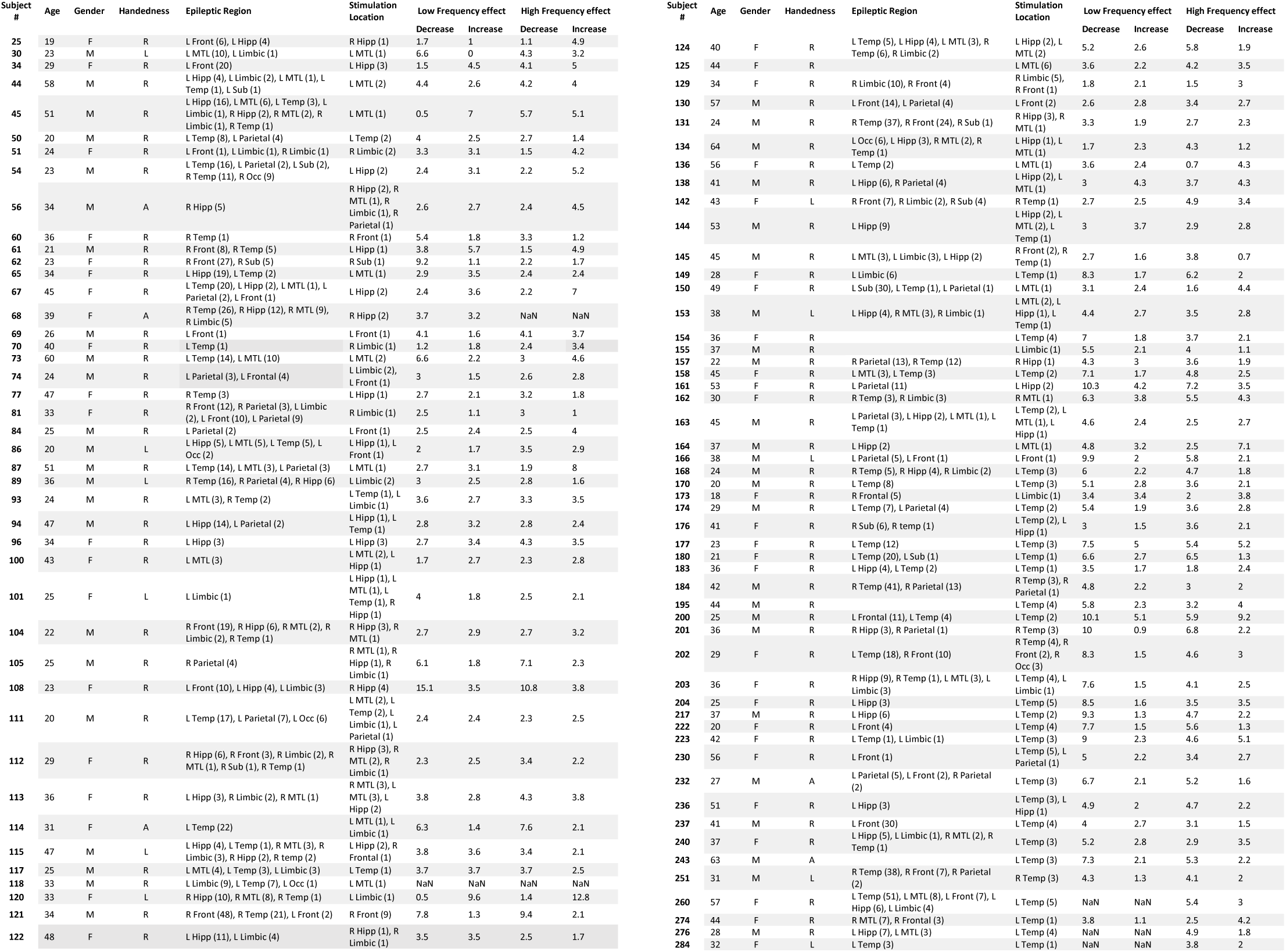
Subject summary table. Gender: M: Male, F: Female; Handedness: R: right, L: left, A: ambidextrous, U: undetermined. Electrode locations: R/L: right/left; Front: frontal cortex; Temp: Temporal cortex; MTL: Medial Temporal Lobe (non-hippocampal); Hipp: hippocampus. Numbers in parentheses indicate number of bipolar contacts in each area for both clinically determined epileptic regions and stimulation sites. Columns labeled “Low-” and “high-frequency effects” indicates the average number of recording electrodes in each subject that show significant HFA increases or decreases, averaged across stimulation sites and amplitudes for low-(10–50 Hz) and high-frequency (100–200 Hz) stimulation, respectively.

**Table S2:**
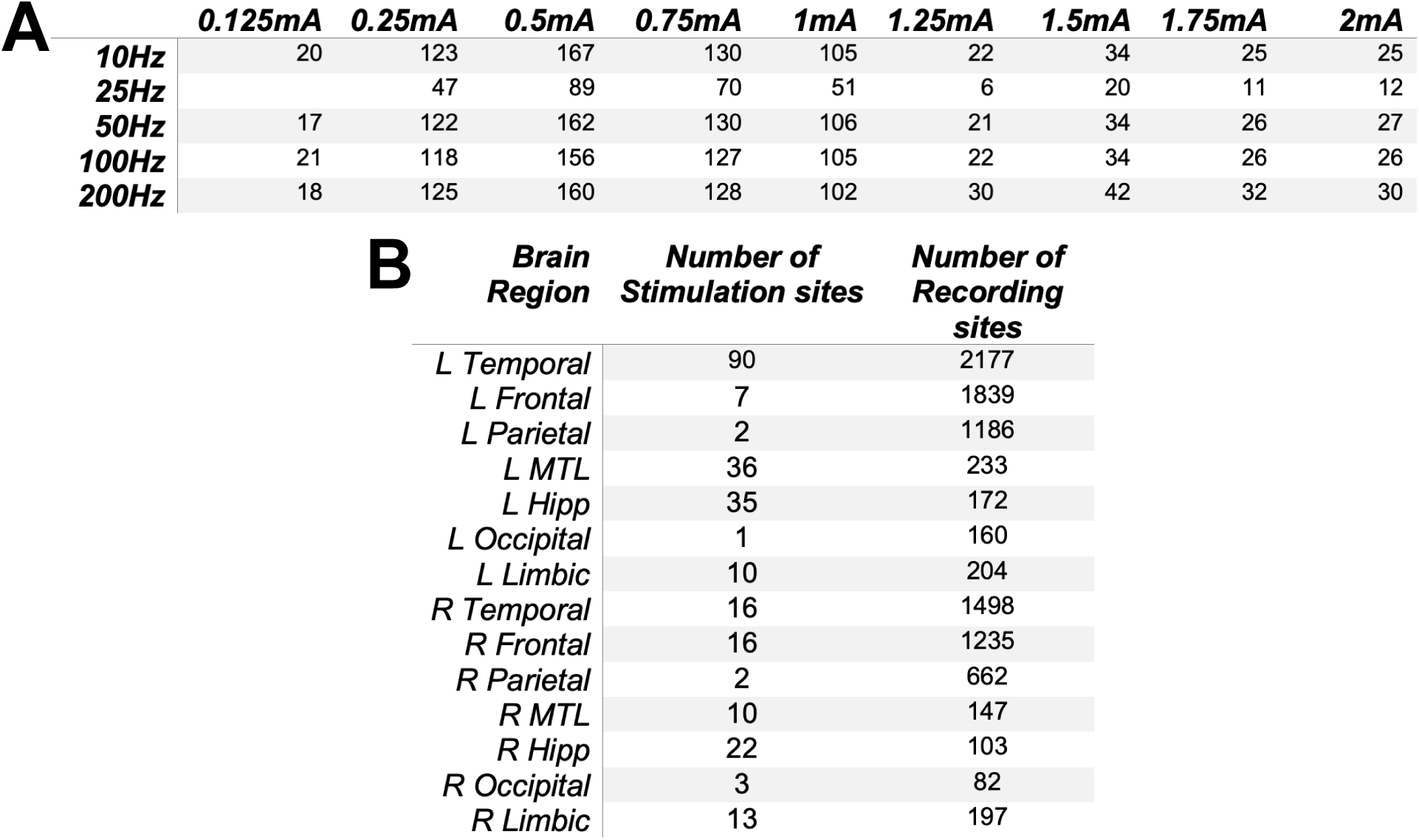
Number of stimulation sites across subjects. **(A)** Number of total stimulation sites used in population analyses across subjects for each combination of frequency and amplitude. **(B)** Number of stimulation and recording sites in each brain region across subjects.

**Table S3:**
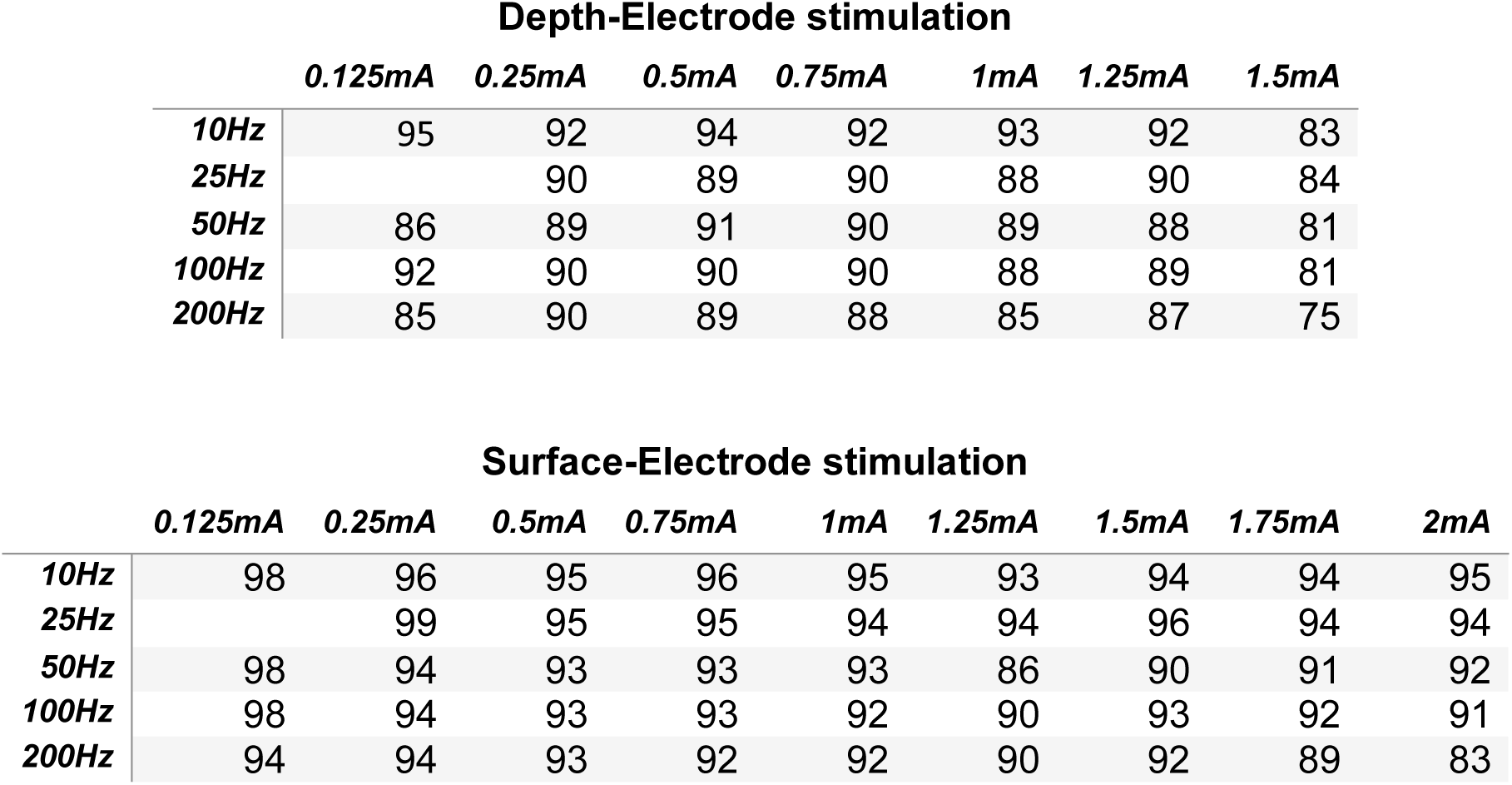
Percent non-artifactual recording sites, by stimulation parameter. Average percent of recording electrodes by stimulation site type (depth-top; surface-bottom), frequency, and amplitude that were included in analyses after being determined as non-artifactual by artifact rejection algorithm.

| Surface-Electrode stimulation |  |  |  |  |  |  |  |  |  |
| --- | --- | --- | --- | --- | --- | --- | --- | --- | --- |
|  | 0.125mA | 0.25mA | 0.5mA | 0.75mA | 1mA | 1.25mA | 1.5mA | 1.75mA | 2mA |
| 10Hz | 98 | 96 | 95 | 96 | 95 | 93 | 94 | 94 | 95 |
| 25Hz |  | 99 | 95 | 95 | 94 | 94 | 96 | 94 | 94 |
| 50Hz | 98 | 94 | 93 | 93 | 93 | 86 | 90 | 91 | 92 |
| 100Hz | 98 | 94 | 93 | 93 | 92 | 90 | 93 | 92 | 91 |
| 200Hz | 94 | 94 | 93 | 92 | 92 | 90 | 92 | 89 | 83 |

**Table S4:**
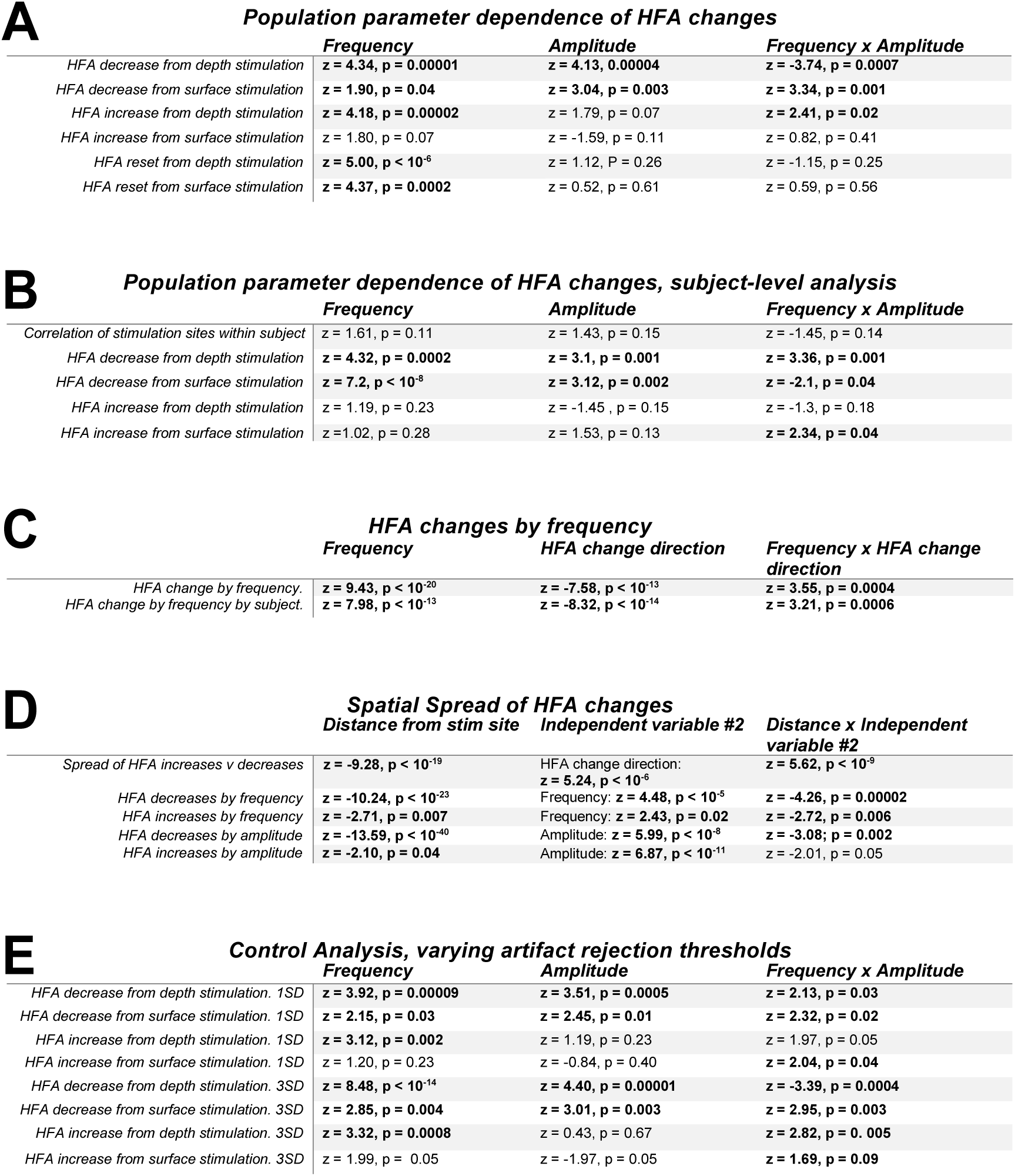
Results of fitted linear mixed effects (LME) models. Each table row indicates one fitted model. Columns indicate model independent parameters. Bold values indicates p < 0.05. **(A)** LME models analyzing the dependence of HFA decreases, increases, and resetting on stimulation frequency and amplitude across the population (Figs. 2A–B and 5C). **(B)** LME models examining subject-level effects of HFA power changes (Fig. S3B–C) and of correlation between HFA changes across stimulation sites (Fig. S3A). **(C)** LME model analysis of HFA decreases and increases by frequency only (Figs. 2C, S3D). **(D)** LME model analysis of the spatial spread of HFA decreases and increases (Fig. 4A–D). Models were separately computed to identify how the spread of HFA changes vary with stimulation frequency (Fig. 4B) and amplitude (Fig. 4C). **(E)** Artifact-rejection control analysis comparing the dependence of HFA decreases and increases on stimulation frequency and amplitude (Fig. S6A,C) with the main population results. Results from artifact rejection thresholds of 1 SD and 3 SD are comparable to those found with the 2-SD threshold, as used in the main analyses (Table S4A).

|  | Frequency | Amplitude | Frequency x Amplitude |
| --- | --- | --- | --- |
| HFA decrease from depth stimulation | **z = 4.34, p = 0.00001** | **z = 4.13, 0.00004** | **z = -3.74, p = 0.0007** |
| HFA decrease from surface stimulation | **z = 1.90, p = 0.04** | **z = 3.04, p = 0.003** | **z = 3.34, p = 0.001** |
| HFA increase from depth stimulation | **z = 4.18, p = 0.00002** | **z = 1.79, p = 0.07** | **z = 2.41, p = 0.02** |
| HFA increase from surface stimulation | **z = 1.80, p = 0.07** | **z = -1.59, p = 0.11** | **z = 0.82, p = 0.41** |
| HFA reset from depth stimulation | **z = 5.00, p < 10<sup>-6</sup>** | **z = 1.12, P = 0.26** | **z = -1.15, p = 0.25** |
| HFA reset from surface stimulation | **z = 4.37, p = 0.0002** | **z = 0.52, p = 0.61** | **z = 0.59, p = 0.56** |

B Population parameter dependence of HFA changes, subject-level analysis
|  | Frequency | Amplitude | Frequency x Amplitude |
| --- | --- | --- | --- |
| Correlation of stimulation sites within subject | **z = 1.61, p = 0.11** | **z = 1.43, p = 0.15** | **z = -1.45, p = 0.14** |
| HFA decrease from depth stimulation | **z = 4.32, p = 0.0002** | **z = 3.1, p = 0.001** | **z = 3.36, p = 0.001** |
| HFA decrease from surface stimulation | **z = 7.2, p < 10<sup>-8</sup>** | **z = 3.12, p = 0.002** | **z = -2.1, p = 0.04** |
| HFA increase from depth stimulation | **z = 1.19, p = 0.23** | **z = -1.45, p = 0.15** | **z = -1.3, p = 0.18** |
| HFA increase from surface stimulation | **z = 1.02, p = 0.28** | **z = 1.53, p = 0.13** | **z = 2.34, p = 0.04** |

C HFA changes by frequency
|  | Frequency | HFA change direction | Frequency x HFA change direction |
| --- | --- | --- | --- |
| HFA change by frequency. | **z = 9.43, p < 10<sup>-20</sup>** | **z = -7.58, p < 10<sup>-13</sup>** | **z = 3.55, p = 0.0004** |
| HFA change by frequency by subject. | **z = 7.98, p < 10<sup>-13</sup>** | **z = -8.32, p < 10<sup>-14</sup>** | **z = 3.21, p = 0.0006** |

D Spatial Spread of HFA changes
|  | Distance from stim site | Independent variable #2 | Distance x Independent variable #2 |
| --- | --- | --- | --- |
| Spread of HFA increases v decreases | **z = -9.28, p < 10<sup>-19</sup>** | HFA change direction:<br>**z = 5.24, p < 10<sup>-6</sup>** | **z = 5.62, p < 10<sup>-9</sup>** |
| HFA decreases by frequency | **z = -10.24, p < 10<sup>-23</sup>** | Frequency: **z = 4.48, p < 10<sup>-5</sup>** | **z = -4.26, p = 0.00002** |
| HFA increases by frequency | **z = -2.71, p = 0.007** | Frequency: **z = 2.43, p = 0.02** | **z = -2.72, p = 0.006** |
| HFA decreases by amplitude | **z = -13.59, p < 10<sup>-40</sup>** | Amplitude: **z = 5.99, p < 10<sup>-8</sup>** | **z = -3.08, p = 0.002** |
| HFA increases by amplitude | **z = -2.10, p = 0.04** | Amplitude: **z = 6.87, p < 10<sup>-11</sup>** | **z = -2.01, p = 0.05** |

**Table S4: Results of fitted linear mixed effects (LME) models.**
|  | Frequency | Amplitude | Frequency x Amplitude |
| --- | --- | --- | --- |
| HFA decrease from depth stimulation. 1SD | **z = 3.92, p = 0.00009** | **z = 3.51, p = 0.0005** | **z = 2.13, p = 0.03** |
| HFA decrease from surface stimulation. 1SD | **z = 2.15, p = 0.03** | **z = 2.45, p = 0.01** | **z = 2.32, p = 0.02** |
| HFA increase from depth stimulation. 1SD | **z = 3.12, p = 0.002** | **z = 1.19, p = 0.23** | **z = 1.97, p = 0.05** |
| HFA increase from surface stimulation. 1SD | **z = 1.20, p = 0.23** | **z = -0.84, p = 0.40** | **z = 2.04, p = 0.04** |
| HFA decrease from depth stimulation. 3SD | **z = 8.48, p < 10<sup>-14</sup>** | **z = 4.40, p = 0.00001** | **z = -3.39, p = 0.0004** |
| HFA decrease from surface stimulation. 3SD | **z = 2.85, p = 0.004** | **z = 3.01, p = 0.003** | **z = 2.95, p = 0.003** |
| HFA increase from depth stimulation. 3SD | **z = 3.32, p = 0.0008** | **z = 0.43, p = 0.67** | **z = 2.82, p = 0.005** |
| HFA increase from surface stimulation. 3SD | **z = 1.99, p = 0.05** | **z = -1.97, p = 0.05** | **z = 1.69, p = 0.09** |

