## Supplemental Analyses for "The effects of direct brain stimulation in humans depend on frequency, amplitude, and white-matter proximity"

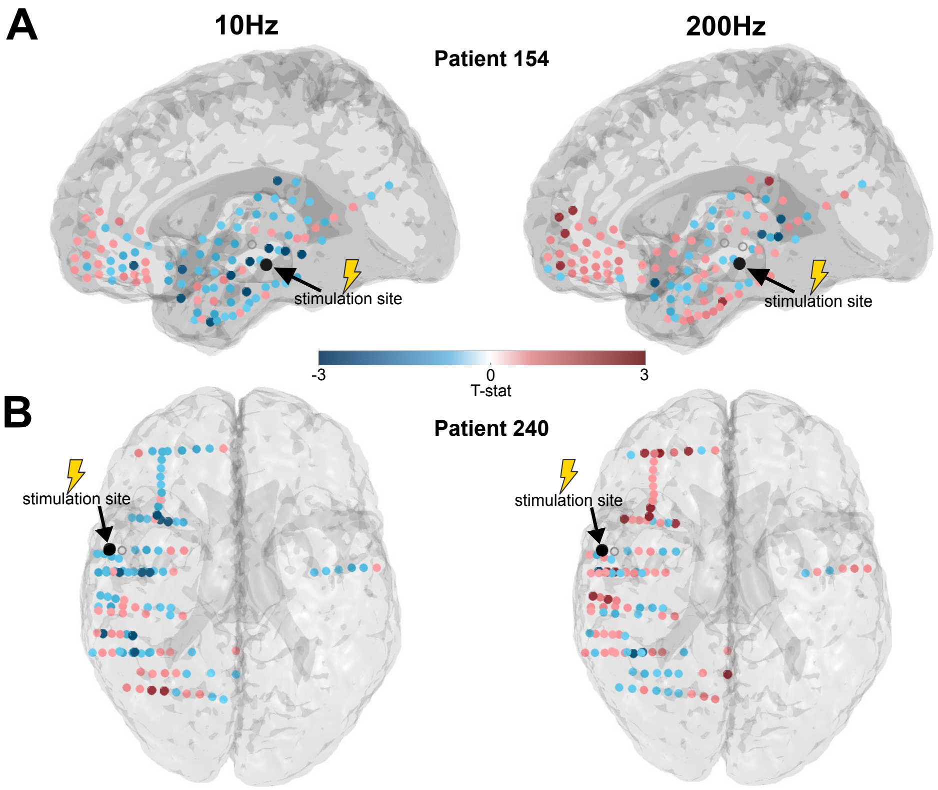


**Figure S1: HFA responses depend on stimulation frequency: Additional example subjects.** **(A)** *Brain maps showing the mean HFA responses across recording electrodes to 10-Hz (left) and 200-Hz (right) stimulation at the same site in Patient 154. The stimulation site is indicated in black and color indicates the t statistic of the change in HFA power from stimulation at each recording electrode. The recording electrodes that are excluded due to artifacts are indicated with an open gray circle.* **(B)** *Brain maps of HFA responses to 10-Hz (left) and 200-Hz (right) stimulation at the same site in example Patient 240. Plot format follows panel A.*

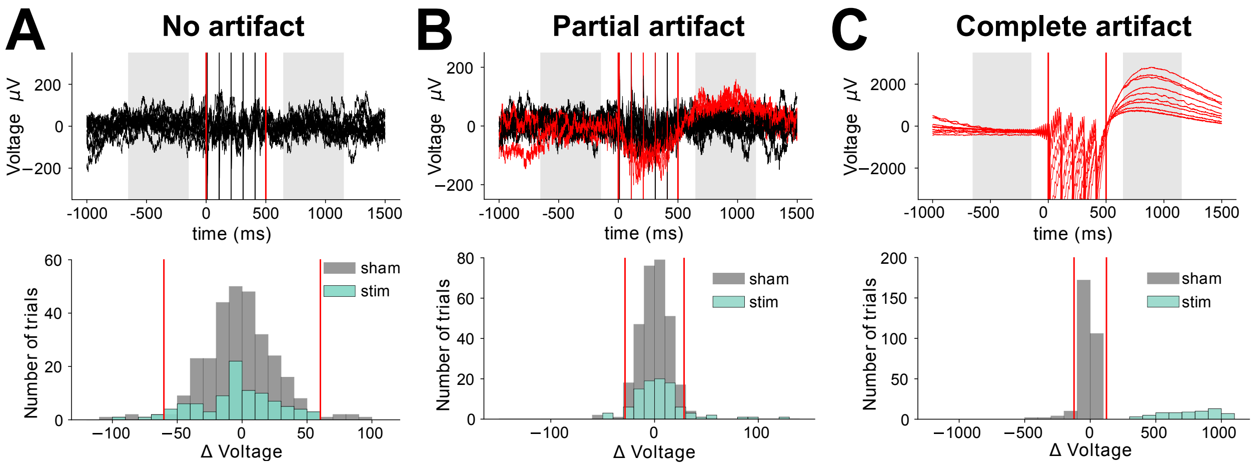


**Figure S2: Illustration of our methods for detecting post-stimulation artifacts.** **(A)** *Top panel, raw signals from 10 trials recorded on an example electrode (Patient 195, electrode 21). Shading indicates the 500-ms time periods before and after each stimulation trial when we measured HFA power. Red lines denote stimulation onset and offset. Bottom panel, histogram of the differences in voltage (post- minus pre-stimulation) for trials when stimulation was applied (turquoise) as well as sham trials (gray). Red lines indicate the artifact-rejection threshold (2SD of sham distribution). Because all of the voltage values on stimulation trials fell within the inner bounds of the thresholds, no trials were rejected.* **(B)** *An analogous plot to Panel A, created from data of 10 trials on a different example recording electrode (Patient 195, electrode 22). Here, two trials (shown in red) were identified as showing post-stimulation artifacts because their change in voltage (POST-PRE) fell outside the 2-SD threshold computed from the voltage difference measured on that electrode for sham trials (see bottom panel).* **(C)** *This plot shows data from a third electrode (Patient 195, electrode 18), where all ten trials were identified as showing post-stimulation artifacts. This entire electrode was excluded from subsequent analyses because all trials showed artifacts (see* Methods*).*

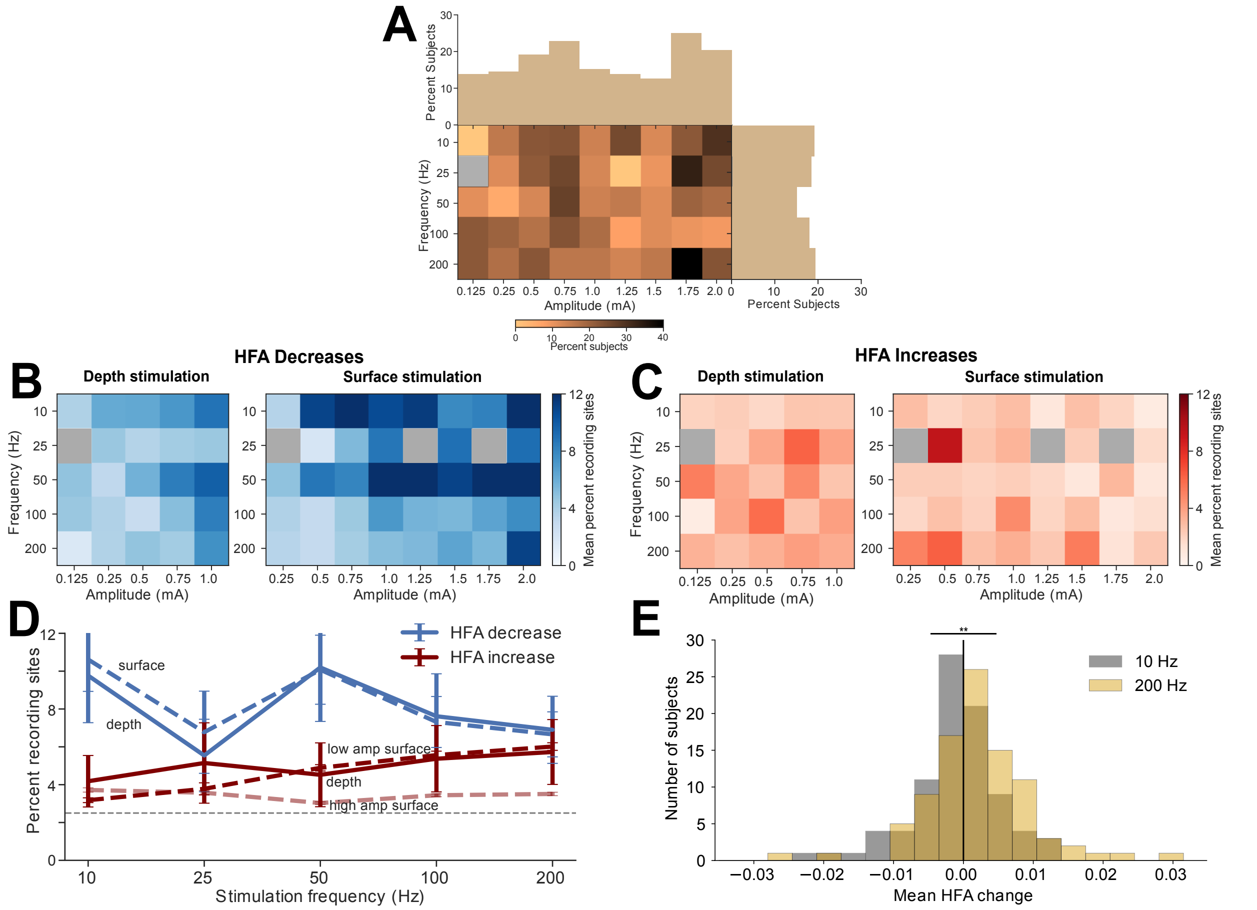


**Figure S3: Subject-level analyses of the effect of stimulation frequency and amplitude on HFA power. (A)** *To test whether a subject showed the same response pattern across different stimulation sites, we computed the intraclass correlation coefficent (ICC) between HFA patterns produced by different stimulation sites. A significant ICC indicates that a similar brain-wide HFA pattern was created by stimulating at different locations in the same subject. This plot illustrates, for each frequency and amplitude, the percentage of subjects that showed similar response patterns across different stimulation sites (as identified with a significant positive ICC). This analysis showed that, on average, 16% of subjects show similar HFA patterns for multiple stimulation sites. Because of this above-chance similarity across stimulation sites, we conducted subject-level analyses of the effects of stimulation, rather than stimulation site-level analyses as in Figure 2.* **(B)** *Subject-level analysis of the mean percent of recording electrodes that showed significant HFA decreases for each combination of stimulation frequency and amplitude, separately computed for depth (left) and surface (right) stimulation. LME modeling shows a similar pattern of statistical effects as in Figure 2A (see Table S4).* **(C)** *Subject-level analysis of the mean percent of recording electrodes that show significant HFA increases for each combination of stimulation parameters. Again, LME modeling shows similar results as Figure 2B.* **(D)** *Subject-level analysis that is analogous to Figure 2C. Direction of HFA change × Frequency interaction: $z = 3.21$; $p = 0.0006$, LME model).* **(E)** *Subject-level analysis that is analogous to Figure 2D. These distributions differ significantly ($z = -3.82$, $p < 10^{-3}$, rank-sum test).*

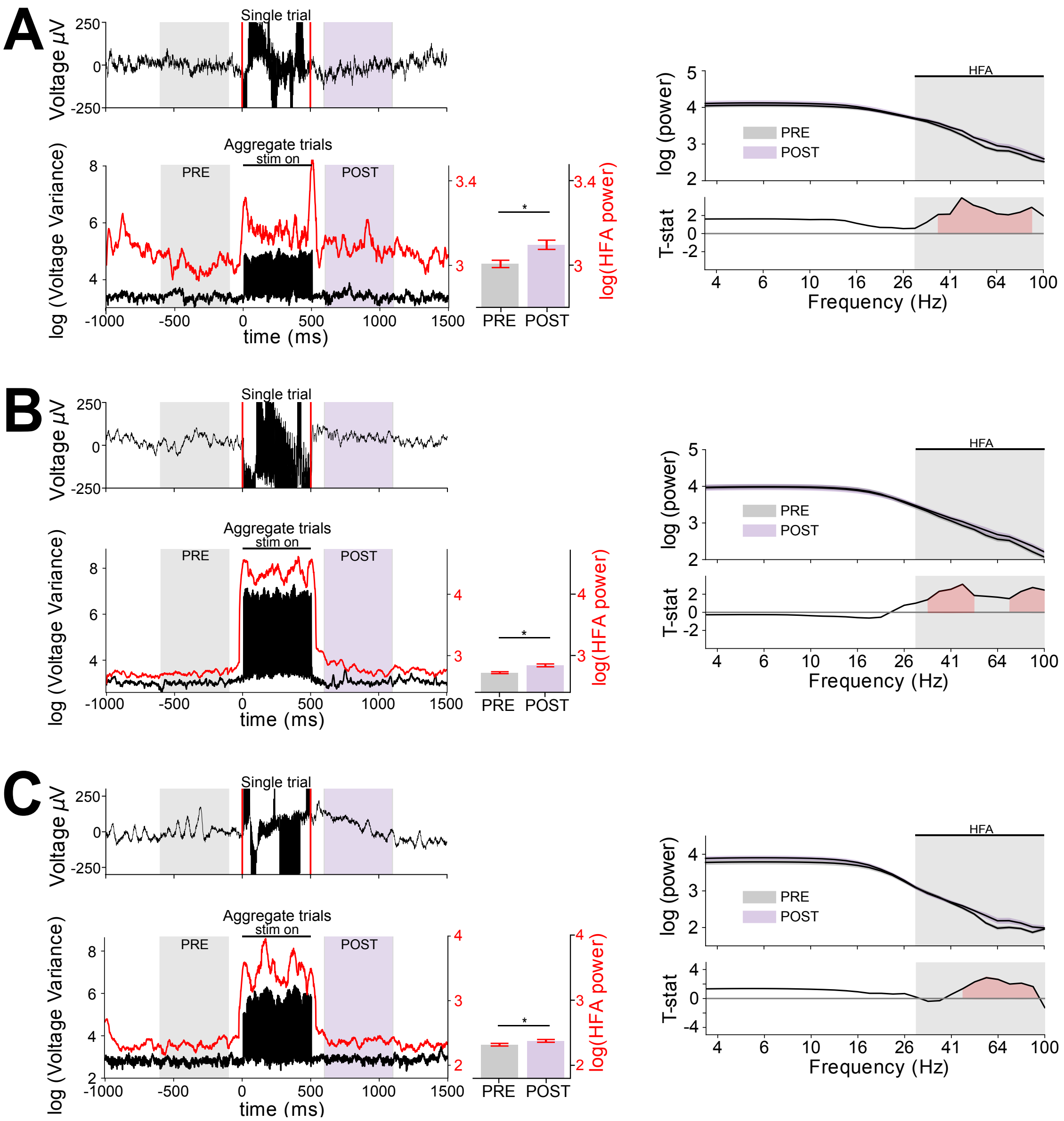


**Figure S4: Illustration of how our analysis method avoids high-frequency stimulation artifacts. (A)** *Top-left panel, raw signals (black line) from one trial recorded from Patient 195, Electrode 67. Red lines denote stimulation onset and offset. Bottom-left panel, illustration of the timecourse of stimulation artifacts seen on this channel. Black line indicates the variance of voltage measurements across trials at each timepoint. The marked increase in variance indicates that stimulation artifact affects recordings specifically when the stimulator was active ("stim on"). The red line indicates the mean HFA power. The gray and purple shading indicates the pre and post-stimulation analysis periods. Critically, as this plot shows, the impact of stimulation artifacts on HFA power consistently drops off before and after stimulation and does not overlap with the pre- (gray) or post-stimulation (purple) analysis periods. Right-top panel, log-transformed mean power spectrum for the pre- and post-stimulation intervals. This plot illustrates that stimulation most strongly increases activity in the HFA band (gray shading from 30-100 Hz). Right-bottom panel, t statistic of the difference in pre- and post-stimulation power at each frequency. Red shading indicates positive significant differences at p<0.05.* **(B)** *Plots follow panel A for Patient 154, Electrode 37* **(A)** *Plots follow panel A for Patient 240, Electrode 36*

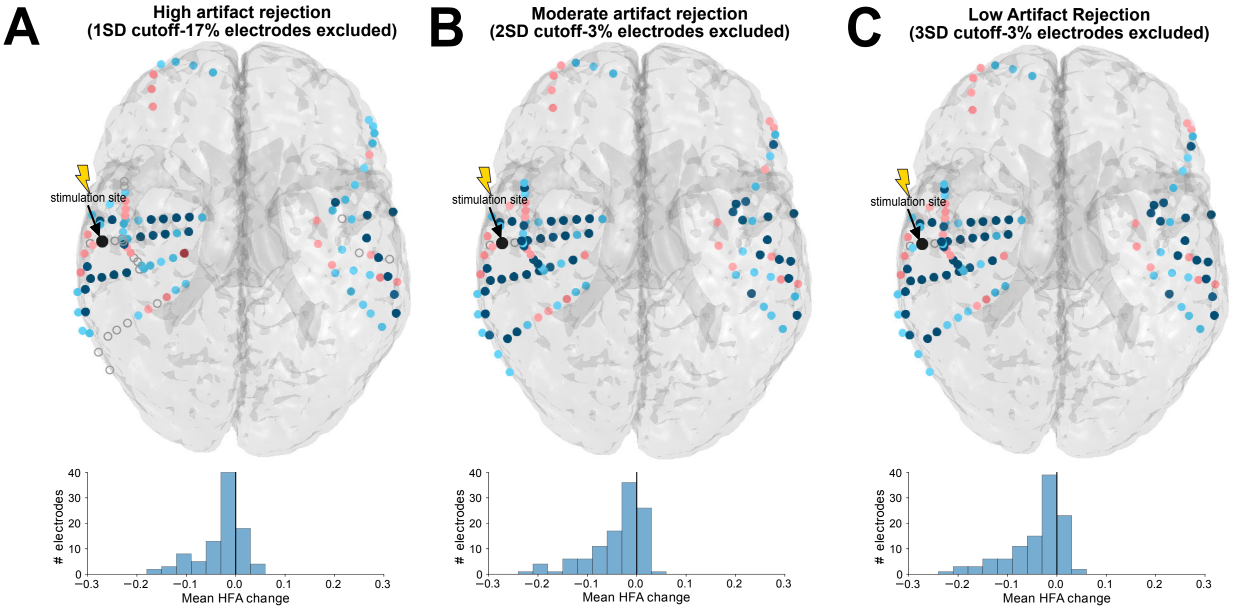


**Figure S5: Effects of different artifact-rejection thresholds on HFA power: Data from one example subject.** **(A)** *Illustration of HFA power changes in patient 195 from 10-Hz, 1-mA stimulation after artifact rejection with a 1-SD cutoff, which excludes 31% of trials and 17% of recording electrodes. Top panel indicates the brain-wide mapping of HFA power changes. Recording electrodes excluded due to artifact indicated are indicated by an open gray circle. Bottom panel, the distribution of mean HFA changes across all analyzed recording electrodes in this subject.* **(B)** *Analysis of HFA power changes in this patient with a 2-SD cutoff, which excludes 11% of trials and 3% of recording electrodes.* **(C)** *Analysis of HFA power changes in this patient with a 3-SD cutoff, which excludes 5% of trials and 3% of recording electrodes.*

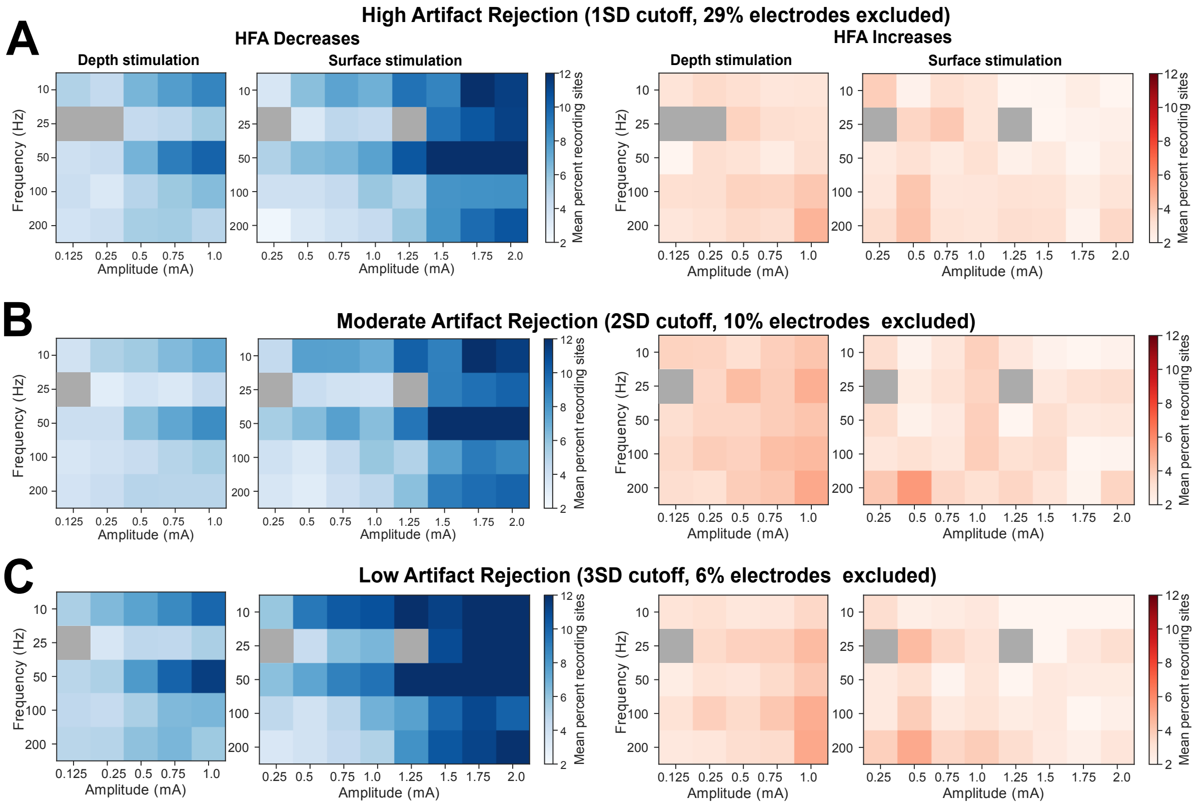


**Figure S6: Effects of different artifact-rejection thresholds on HFA power: Population-level analysis. (A)** *Analysis of the percent of recording sites where HFA significantly increased or decreased for when using a 1-SD artifact-rejection threshold. With this threshold we excluded 29% of recording electrodes and 28% of stimulation trials on the remaining electrodes. LME model analysis confirmed a similar relationship between HFA changes and stimulation parameters as in Figure 2A,B (see Table S4).* **(B)** *Same analysis as our main population results in Figure 2 using a 2-SD artifact-rejection threshold. With this threshold, we excluded 10% of recording electrodes and 12% of trials on the remaining electrodes.* **(C)** *Same analysis as above, but using a 3-SD artifact-rejection threshold. Here, we excluded 6% of recording electrodes and 5% of trials on remaining electrodes. LME model analysis confirmed a similar relationship between HFA changes and stimulation parameters as in Figure 2A,B (see Table S4).*

| Subject # | Age | Gender | Handedness | Epileptic Region | Stimulation Location | Low Frequency effect | | High Frequency effect | |
|---|---|---|---|---|---|---|---|---|---|
| | | | | | | Decrease | Increase | Decrease | Increase |
| **25** | 19 | F | R | L Front (6), L Hipp (4) | R Hipp (1) | 1.7 | 1 | 1.1 | 4.9 |
| **30** | 23 | M | L | L MTL (10), L Limbic (1) | L MTL (1) | 6.6 | 0 | 4.3 | 3.2 |
| **34** | 29 | F | R | L Front (20) | L Hipp (3) | 1.5 | 4.5 | 4.1 | 5 |
| **44** | 58 | M | R | L Hipp (4), L Limbic (2), L MTL (1), L Temp (1), L Sub (1) | L MTL (2) | 4.4 | 2.6 | 4.2 | 4 |
| **45** | 51 | M | R | L Hipp (16), L MTL (6), L Temp (3), L Limbic (1), R Hipp (2), R MTL (2), R Limbic (1), R Temp (1) | L MTL (1) | 0.5 | 7 | 5.7 | 5.1 |
| **50** | 20 | M | R | L Temp (8), L Parietal (4) | L Temp (2) | 4 | 2.5 | 2.7 | 1.4 |
| **51** | 24 | F | R | L Front (1), L Limbic (1), R Limbic (1) | R Limbic (2) | 3.3 | 3.1 | 1.5 | 4.2 |
| **54** | 23 | M | R | L Temp (16), L Parietal (2), L Sub (2), R Temp (11), R Occ (9) | L Hipp (2) | 2.4 | 3.1 | 2.2 | 5.2 |
| **56** | 34 | M | A | R Hipp (5) | R Hipp (2), R MTL (1), R Limbic (1), R Parietal (1) | 2.6 | 2.7 | 2.4 | 4.5 |
| **60** | 36 | F | R | R Temp (1) | R Front (1) | 5.4 | 1.8 | 3.3 | 1.2 |
| **61** | 21 | M | R | R Front (8), R Temp (5) | L Hipp (1) | 3.8 | 5.7 | 1.5 | 4.9 |
| **62** | 23 | F | R | R Front (27), R Sub (5) | R Sub (1) | 9.2 | 1.1 | 2.2 | 1.7 |
| **65** | 34 | F | R | L Hipp (19), L Temp (2) | L MTL (1) | 2.9 | 3.5 | 2.4 | 2.4 |
| **67** | 45 | F | R | L Temp (20), L Hipp (2), L MTL (1), L Parietal (2), L Front (1) | L Hipp (2) | 2.4 | 3.6 | 2.2 | 7 |
| **68** | 39 | F | A | R Temp (26), R Hipp (12), R MTL (9), R Limbic (5) | R Hipp (2) | 3.7 | 3.2 | NaN | NaN |
| **69** | 26 | M | R | L Front (1) | L Front (1) | 4.1 | 1.6 | 4.1 | 3.7 |
| **70** | 40 | F | R | L Temp (1) | R Limbic (1) | 1.2 | 1.8 | 2.4 | 3.4 |
| **73** | 60 | M | R | L Temp (14), L MTL (10) | L MTL (2) | 6.6 | 2.2 | 3 | 4.6 |
| **74** | 24 | M | R | L Parietal (3), L Frontal (4) | L Limbic (2), L Front (1) | 3 | 1.5 | 2.6 | 2.8 |
| **77** | 47 | F | R | R Temp (3) | L Hipp (1) | 2.7 | 2.1 | 3.2 | 1.8 |
| **81** | 33 | F | R | R Front (12), R Parietal (3), L Limbic (2), L Front (10), L Parietal (9) | R Limbic (1) | 2.5 | 1.1 | 3 | 1 |
| **84** | 25 | M | R | L Parietal (2) | L Front (1) | 2.5 | 2.4 | 2.5 | 4 |
| **86** | 20 | M | L | L Hipp (5), L MTL (5), L Temp (5), L Occ (2) | L Hipp (1), L Front (1) | 2 | 1.7 | 3.5 | 2.9 |
| **87** | 51 | M | R | L Temp (14), L MTL (3), L Parietal (3) | L MTL (1) | 2.7 | 3.1 | 1.9 | 8 |
| **89** | 36 | M | L | R Temp (16), R Parietal (4), R Hipp (6) | L Limbic (2) | 3 | 2.5 | 2.8 | 1.6 |
| **93** | 24 | M | R | L MTL (3), R Temp (2) | L Temp (1), L Limbic (1) | 3.6 | 2.7 | 3.3 | 3.5 |
| **94** | 47 | M | R | L Hipp (14), L Parietal (2) | L Hipp (1), L Temp (1) | 2.8 | 3.2 | 2.8 | 2.4 |
| **96** | 34 | F | R | L Hipp (3) | L Hipp (3) | 2.7 | 3.4 | 4.3 | 3.5 |
| **100** | 43 | F | R | L MTL (3) | L MTL (2), L Hipp (1) | 1.7 | 2.7 | 2.3 | 2.8 |
| **101** | 25 | F | L | L Limbic (1) | L Hipp (1), L MTL (1), L Temp (1), R Hipp (1) | 4 | 1.8 | 2.5 | 2.1 |
| **104** | 22 | M | R | R Front (19), R Hipp (6), R MTL (2), R Limbic (2), R Temp (1) | R Hipp (3), R MTL (1) | 2.7 | 2.9 | 2.7 | 3.2 |
| **105** | 25 | M | R | R Parietal (4) | R MTL (1), R Hipp (1), R Limbic (1) | 6.1 | 1.8 | 7.1 | 2.3 |
| **108** | 23 | F | R | L Front (10), L Hipp (4), L Limbic (3) | R Hipp (4) | 15.1 | 3.5 | 10.8 | 3.8 |
| **111** | 20 | M | R | L Temp (17), L Parietal (7), L Occ (6) | L MTL (2), L Temp (2), L Limbic (1), L Parietal (1) | 2.4 | 2.4 | 2.3 | 2.5 |
| **112** | 29 | F | R | R Hipp (6), R Front (3), R Limbic (2), R MTL (1), R Sub (1), R Temp (1) | R Hipp (3), R MTL (2), R Limbic (1) | 2.3 | 2.5 | 3.4 | 2.2 |
| **113** | 36 | F | R | L Hipp (3), R Limbic (2), R MTL (1) | R MTL (3), L MTL (3), L Hipp (2) | 3.8 | 2.8 | 4.3 | 3.8 |
| **114** | 31 | F | A | L Temp (22) | L MTL (1), L Limbic (1) | 6.3 | 1.4 | 7.6 | 2.1 |
| **115** | 47 | M | L | L Hipp (4), L Temp (1), R MTL (3), R Limbic (3), R Hipp (2), R temp (2) | L Hipp (2), R Frontal (1) | 3.8 | 3.6 | 3.4 | 2.1 |
| **117** | 25 | M | R | L MTL (4), L Temp (3), L Limbic (3) | L Temp (1) | 3.7 | 3.7 | 3.7 | 2.5 |
| **118** | 33 | M | R | L Limbic (9), L Temp (7), L Occ (1) | L MTL (1) | NaN | NaN | NaN | NaN |
| **120** | 33 | F | L | R Hipp (10), R MTL (8), R Temp (1) | L Limbic (1) | 0.5 | 9.6 | 1.4 | 12.8 |
| **121** | 34 | M | R | R Front (48), R Temp (21), L Front (2) | R Front (9) | 7.8 | 1.3 | 9.4 | 2.1 |
| **122** | 48 | F | R | L Hipp (11), L Limbic (4) | R Hipp (1), R Limbic (1) | 3.5 | 3.5 | 2.5 | 1.7 |
| **124** | 40 | F | R | L Temp (5), L Hipp (4), L MTL (3), R Temp (6), R Limbic (2) | L Hipp (2), L MTL (2) | 5.2 | 2.6 | 5.8 | 1.9 |
| **125** | 44 | F | R | | L MTL (6) | 3.6 | 2.2 | 4.2 | 3.5 |
| **129** | 34 | F | R | R Limbic (10), R Front (4) | R Limbic (5), R Front (1) | 1.8 | 2.1 | 1.5 | 3 |
| **130** | 57 | M | R | L Front (14), L Parietal (4) | L Front (2) | 2.6 | 2.8 | 3.4 | 2.7 |
| **131** | 24 | M | R | R Temp (37), R Front (24), R Sub (1) | R Hipp (3), R MTL (1) | 3.3 | 1.9 | 2.7 | 2.3 |
| **134** | 64 | M | R | L Occ (6), L Hipp (3), R MTL (2), R Temp (1) | L Hipp (1), L MTL (1) | 1.7 | 2.3 | 4.3 | 1.2 |
| **136** | 56 | F | R | L Temp (2) | L MTL (1) | 3.6 | 2.4 | 0.7 | 4.3 |
| **138** | 41 | M | R | L Hipp (6), R Parietal (4) | L Hipp (2), L MTL (1) | 3 | 4.3 | 3.7 | 4.3 |
| **142** | 43 | F | L | R Front (7), R Limbic (2), R Sub (4) | R Temp (1) | 2.7 | 2.5 | 4.9 | 3.4 |
| **144** | 53 | M | R | L Hipp (9) | L Hipp (2), L MTL (2), L Temp (1) | 3 | 3.7 | 2.9 | 2.8 |
| **145** | 45 | M | R | L MTL (3), L Limbic (3), L Hipp (2) | R Front (2), R Temp (1) | 2.7 | 1.6 | 3.8 | 0.7 |
| **149** | 28 | F | R | L Limbic (6) | L Temp (1) | 8.3 | 1.7 | 6.2 | 2 |
| **150** | 49 | F | R | L Sub (30), L Temp (1), L Parietal (1) | L MTL (1) | 3.1 | 2.4 | 1.6 | 4.4 |
| **153** | 38 | M | L | L Hipp (4), R MTL (3), R Limbic (1) | L MTL (2), L Hipp (1), L Temp (1) | 4.4 | 2.7 | 3.5 | 2.8 |
| **154** | 36 | F | R | | L Temp (4) | 7 | 1.8 | 3.7 | 2.1 |
| **155** | 37 | M | R | | L Limbic (1) | 5.5 | 2.1 | 4 | 1.1 |
| **157** | 22 | M | R | R Parietal (13), R Temp (12) | R Hipp (1) | 4.3 | 3 | 3.6 | 1.9 |
| **158** | 45 | F | R | L MTL (3), L Temp (3) | L Temp (2) | 7.1 | 1.7 | 4.8 | 2.5 |
| **161** | 53 | F | R | L Parietal (11) | L Hipp (2) | 10.3 | 4.2 | 7.2 | 3.5 |
| **162** | 30 | F | R | R Temp (3), R Limbic (3) | R MTL (1) | 6.3 | 3.8 | 5.5 | 4.3 |
| **163** | 45 | M | R | L Parietal (3), L Hipp (2), L MTL (1), L Temp (1) | L Temp (2), L MTL (1), L Hipp (1) | 4.6 | 2.4 | 2.5 | 2.7 |
| **164** | 37 | M | R | L Hipp (2) | L MTL (1) | 4.8 | 3.2 | 2.5 | 7.1 |
| **166** | 38 | M | L | L Parietal (5), L Front (1) | L Front (1) | 9.9 | 2 | 5.8 | 2.1 |
| **168** | 24 | M | R | R Temp (5), R Hipp (4), R Limbic (2) | L Temp (3) | 6 | 2.2 | 4.7 | 1.8 |
| **170** | 20 | M | R | L Temp (8) | L Temp (3) | 5.1 | 2.8 | 3.6 | 2.1 |
| **173** | 18 | F | R | R Frontal (5) | L Limbic (1) | 3.4 | 3.4 | 2 | 3.8 |
| **174** | 29 | M | R | L Temp (7), L Parietal (4) | L Temp (2) | 5.4 | 1.9 | 3.6 | 2.8 |
| **176** | 41 | F | R | R Sub (6), R temp (1) | L Temp (2), L Hipp (1) | 3 | 1.5 | 3.6 | 2.1 |
| **177** | 23 | F | R | L Temp (12) | L Temp (3) | 7.5 | 5 | 5.4 | 5.2 |
| **180** | 21 | F | R | L Temp (20), L Sub (1) | L Temp (1) | 6.6 | 2.7 | 6.5 | 1.3 |
| **183** | 36 | F | R | L Hipp (4), L Temp (2) | L Temp (1) | 3.5 | 1.7 | 1.8 | 2.4 |
| **184** | 42 | M | R | R Temp (41), R Parietal (13) | R Temp (3), R Parietal (1) | 4.8 | 2.2 | 3 | 2 |
| **195** | 44 | M | R | | L Temp (4) | 5.8 | 2.3 | 3.2 | 4 |
| **200** | 25 | M | R | L Frontal (11), L Temp (4) | L Temp (2) | 10.1 | 5.1 | 5.9 | 9.2 |
| **201** | 36 | M | R | R Hipp (3), R Parietal (1) | R Temp (3) | 10 | 0.9 | 6.8 | 2.2 |
| **202** | 29 | F | R | L Temp (18), R Front (10) | R Temp (4), R Front (2), R Occ (3) | 8.3 | 1.5 | 4.6 | 3 |
| **203** | 36 | F | R | R Hipp (9), R Temp (1), L MTL (3), L Limbic (3) | L Temp (4), L Limbic (1) | 7.6 | 1.5 | 4.1 | 2.5 |
| **204** | 25 | F | R | L Hipp (3) | L Temp (5) | 8.5 | 1.6 | 3.5 | 3.5 |
| **217** | 37 | M | R | L Hipp (6) | L Temp (2) | 9.3 | 1.3 | 4.7 | 2.2 |
| **222** | 20 | F | R | L Front (4) | L Temp (4) | 7.7 | 1.5 | 5.6 | 1.3 |
| **223** | 42 | F | R | L Temp (1), L Limbic (1) | L Temp (3) | 9 | 2.3 | 4.6 | 5.1 |
| **230** | 56 | F | R | L Front (1) | L Temp (5), L Parietal (1) | 5 | 2.2 | 3.4 | 2.7 |
| **232** | 27 | M | A | L Parietal (5), L Front (2), R Parietal (2) | L Temp (3) | 6.7 | 2.1 | 5.2 | 1.6 |
| **236** | 51 | F | R | L Hipp (3) | L Temp (3), L Hipp (1) | 4.9 | 2 | 4.7 | 2.2 |
| **237** | 41 | M | R | L Front (30) | L Temp (4) | 4 | 2.7 | 3.1 | 1.5 |
| **240** | 37 | F | R | L Hipp (5), L Limbic (1), R MTL (2), R Temp (1) | L Temp (3) | 5.2 | 2.8 | 2.9 | 3.5 |
| **243** | 63 | M | A | | L Temp (3) | 7.3 | 2.1 | 5.3 | 2.2 |
| **251** | 31 | M | L | R Temp (38), R Front (7), R Parietal (2) | R Temp (3) | 4.3 | 1.3 | 4.1 | 2 |
| **260** | 57 | F | R | L Temp (51), L MTL (8), L Front (7), L Hipp (6), L Limbic (4) | L Temp (5) | NaN | NaN | 5.4 | 3 |
| **274** | 44 | F | R | R MTL (7), R Frontal (3) | L Temp (1) | 3.8 | 1.1 | 2.5 | 4.2 |
| **276** | 28 | M | R | L Hipp (7), L MTL (3) | L Temp (4) | NaN | NaN | 4.9 | 1.8 |
| **284** | 32 | F | L | L Temp (3) | L Temp (1) | NaN | NaN | 3.8 | 2 |

**Table S1: Subject summary table.** *Gender: M: Male, F: Female; Handedness: R: right, L: left, A: ambidextrous, U: undetermined. Electrode locations: R/L: right/left; Front: frontal cortex; Temp: Temporal cortex; MTL: Medial Temporal Lobe (non-hippocampal); Hipp: hippocampus. Numbers in parentheses indicate number of bipolar contacts in each area for both clinically determined epileptic regions and stimulation sites. Columns labeled "Low-" and "high-frequency effects" indicates the average number of recording electrodes in each subject that show significant HFA increases or decreases, averaged across stimulation sites and amplitudes for low- (10–50 Hz) and high-frequency (100–200 Hz) stimulation, respectively.*

**A**

| | 0.125mA | 0.25mA | 0.5mA | 0.75mA | 1mA | 1.25mA | 1.5mA | 1.75mA | 2mA |
|---|---|---|---|---|---|---|---|---|---|
| 10Hz | 20 | 123 | 167 | 130 | 105 | 22 | 34 | 25 | 25 |
| 25Hz | | 47 | 89 | 70 | 51 | 6 | 20 | 11 | 12 |
| 50Hz | 17 | 122 | 162 | 130 | 106 | 21 | 34 | 26 | 27 |
| 100Hz | 21 | 118 | 156 | 127 | 105 | 22 | 34 | 26 | 26 |
| 200Hz | 18 | 125 | 160 | 128 | 102 | 30 | 42 | 32 | 30 |

**B**

| Brain Region | Number of Stimulation sites | Number of Recording sites |
|---|---|---|
| L Temporal | 90 | 2177 |
| L Frontal | 7 | 1839 |
| L Parietal | 2 | 1186 |
| L MTL | 36 | 233 |
| L Hipp | 35 | 172 |
| L Occipital | 1 | 160 |
| L Limbic | 10 | 204 |
| R Temporal | 16 | 1498 |
| R Frontal | 16 | 1235 |
| R Parietal | 2 | 662 |
| R MTL | 10 | 147 |
| R Hipp | 22 | 103 |
| R Occipital | 3 | 82 |
| R Limbic | 13 | 197 |

**Table S2: Number of stimulation sites across subjects. (A)** *Number of total stimulation sites used in population analyses across subjects for each combination of frequency and amplitude.* **(B)** *Number of stimulation and recording sites in each brain region across subjects.*

### Depth-Electrode stimulation

| | 0.125mA | 0.25mA | 0.5mA | 0.75mA | 1mA | 1.25mA | 1.5mA |
|---|---|---|---|---|---|---|---|
| **10Hz** | 95 | 92 | 94 | 92 | 93 | 92 | 83 |
| **25Hz** | | 90 | 89 | 90 | 88 | 90 | 84 |
| **50Hz** | 86 | 89 | 91 | 90 | 89 | 88 | 81 |
| **100Hz** | 92 | 90 | 90 | 90 | 88 | 89 | 81 |
| **200Hz** | 85 | 90 | 89 | 88 | 85 | 87 | 75 |

### Surface-Electrode stimulation

| | 0.125mA | 0.25mA | 0.5mA | 0.75mA | 1mA | 1.25mA | 1.5mA | 1.75mA | 2mA |
|---|---|---|---|---|---|---|---|---|---|
| **10Hz** | 98 | 96 | 95 | 96 | 95 | 93 | 94 | 94 | 95 |
| **25Hz** | | 99 | 95 | 95 | 94 | 94 | 96 | 94 | 94 |
| **50Hz** | 98 | 94 | 93 | 93 | 93 | 86 | 90 | 91 | 92 |
| **100Hz** | 98 | 94 | 93 | 93 | 92 | 90 | 93 | 92 | 91 |
| **200Hz** | 94 | 94 | 93 | 92 | 92 | 90 | 92 | 89 | 83 |

**Table S3: Percent non-artifactual recording sites, by stimulation parameter.** *Average percent of recording electrodes by stimulation site type (depth-top; surface-bottom), frequency, and amplitude that were included in analyses after being determined as non-artifactual by artifact rejection algorithm.*

**A** 

#### *Population parameter dependence of HFA changes*

| | *Frequency* | *Amplitude* | *Frequency x Amplitude* |
|---|---|---|---|
| *HFA decrease from depth stimulation* | **z = 4.34, p = 0.00001** | **z = 4.13, 0.00004** | **z = -3.74, p = 0.0007** |
| *HFA decrease from surface stimulation* | **z = 1.90, p = 0.04** | **z = 3.04, p = 0.003** | **z = 3.34, p = 0.001** |
| *HFA increase from depth stimulation* | **z = 4.18, p = 0.00002** | z = 1.79, p = 0.07 | **z = 2.41, p = 0.02** |
| *HFA increase from surface stimulation* | z = 1.80, p = 0.07 | z = -1.59, p = 0.11 | z = 0.82, p = 0.41 |
| *HFA reset from depth stimulation* | **z = 5.00, p < $10^{-6}$** | z = 1.12, P = 0.26 | z = -1.15, p = 0.25 |
| *HFA reset from surface stimulation* | **z = 4.37, p = 0.0002** | z = 0.52, p = 0.61 | z = 0.59, p = 0.56 |

**B** 

#### *Population parameter dependence of HFA changes, subject-level analysis*

| | *Frequency* | *Amplitude* | *Frequency x Amplitude* |
|---|---|---|---|
| *Correlation of stimulation sites within subject* | z = 1.61, p = 0.11 | z = 1.43, p = 0.15 | z = -1.45, p = 0.14 |
| *HFA decrease from depth stimulation* | **z = 4.32, p = 0.0002** | **z = 3.1, p = 0.001** | **z = 3.36, p = 0.001** |
| *HFA decrease from surface stimulation* | **z = 7.2, p < $10^{-8}$** | **z = 3.12, p = 0.002** | **z = -2.1, p = 0.04** |
| *HFA increase from depth stimulation* | z = 1.19, p = 0.23 | z = -1.45 , p = 0.15 | z = -1.3, p = 0.18 |
| *HFA increase from surface stimulation* | z =1.02, p = 0.28 | z = 1.53, p = 0.13 | **z = 2.34, p = 0.04** |

**C** 

#### *HFA changes by frequency*

| | *Frequency* | *HFA change direction* | *Frequency x HFA change direction* |
|---|---|---|---|
| *HFA change by frequency.* | **z = 9.43, p < $10^{-20}$** | **z = -7.58, p < $10^{-13}$** | **z = 3.55, p = 0.0004** |
| *HFA change by frequency by subject.* | **z = 7.98, p < $10^{-13}$** | **z = -8.32, p < $10^{-14}$** | **z = 3.21, p = 0.0006** |

**D** 

#### *Spatial Spread of HFA changes*

| | *Distance from stim site* | *Independent variable #2* | *Distance x Independent variable #2* |
|---|---|---|---|
| *Spread of HFA increases v decreases* | **z = -9.28, p < $10^{-19}$** | HFA change direction: **z = 5.24, p < $10^{-6}$** | **z = 5.62, p < $10^{-9}$** |
| *HFA decreases by frequency* | **z = -10.24, p < $10^{-23}$** | Frequency: **z = 4.48, p < $10^{-5}$** | **z = -4.26, p = 0.00002** |
| *HFA increases by frequency* | **z = -2.71, p = 0.007** | Frequency: **z = 2.43, p = 0.02** | **z = -2.72, p = 0.006** |
| *HFA decreases by amplitude* | **z = -13.59, p < $10^{-40}$** | Amplitude: **z = 5.99, p < $10^{-8}$** | **z = -3.08; p = 0.002** |
| *HFA increases by amplitude* | **z = -2.10, p = 0.04** | Amplitude: **z = 6.87, p < $10^{-11}$** | z = -2.01, p = 0.05 |

**E** 

#### *Control Analysis, varying artifact rejection thresholds*

| | *Frequency* | *Amplitude* | *Frequency x Amplitude* |
|---|---|---|---|
| *HFA decrease from depth stimulation. 1SD* | **z = 3.92, p = 0.00009** | **z = 3.51, p = 0.0005** | **z = 2.13, p = 0.03** |
| *HFA decrease from surface stimulation. 1SD* | **z = 2.15, p = 0.03** | **z = 2.45, p = 0.01** | **z = 2.32, p = 0.02** |
| *HFA increase from depth stimulation. 1SD* | **z = 3.12, p = 0.002** | z = 1.19, p = 0.23 | z = 1.97, p = 0.05 |
| *HFA increase from surface stimulation. 1SD* | z = 1.20, p = 0.23 | z = -0.84, p = 0.40 | **z = 2.04, p = 0.04** |
| *HFA decrease from depth stimulation. 3SD* | **z = 8.48, p < $10^{-14}$** | **z = 4.40, p = 0.00001** | **z = -3.39, p = 0.0004** |
| *HFA decrease from surface stimulation. 3SD* | **z = 2.85, p = 0.004** | **z = 3.01, p = 0.003** | **z = 2.95, p = 0.003** |
| *HFA increase from depth stimulation. 3SD* | **z = 3.32, p = 0.0008** | z = 0.43, p = 0.67 | **z = 2.82, p = 0. 005** |
| *HFA increase from surface stimulation. 3SD* | z = 1.99, p = 0.05 | z = -1.97, p = 0.05 | **z = 1.69, p = 0.09** |

**Table S4: Results of fitted linear mixed effects (LME) models.** *Each table row indicates one fitted model. Columns indicate model independent parameters. Bold values indicates* $p < 0.05$. **(A)** *LME models analyzing the dependence of HFA decreases, increases, and resetting on stimulation frequency and amplitude across the population (Figs. 2A–B and 5C).* **(B)** *LME models examining subject-level effects of HFA power changes (Fig. S3B–C) and of correlation between HFA changes across stimulation sites (Fig. S3A).* **(C)** *LME model analysis of HFA decreases and increases by frequency only (Figs. 2C, S3D).* **(D)** *LME model analysis of the spatial spread of HFA decreases and increases (Fig. 4A–D). Models were separately computed to identify how the spread of HFA changes vary with stimulation frequency (Fig. 4B) and amplitude (Fig. 4C).* **(E)** *Artifact-rejection control analysis comparing the dependence of HFA decreases and increases on stimulation frequency and amplitude (Fig. S6A,C) with the main population results. Results from artifact rejection thresholds of 1 SD and 3 SD are comparable to those found with the 2-SD threshold, as used in the main analyses (Table S4A).*
